# Selective preservation of fucose-rich oligosaccharides in the North Atlantic Ocean

**DOI:** 10.1101/2024.09.20.613644

**Authors:** Margot Bligh, Hagen Buck-Wiese, Andreas Sichert, Sarah K. Bercovici, Inga Hellige, Hannah Marchant, Morten Iversen, Uwe Sauer, Thorsten Dittmar, Carol Arnosti, Manuel Liebeke, Jan-Hendrik Hehemann

## Abstract

The ocean has a substantial capacity to store carbon dioxide fixed via photosynthesis in dissolved organic molecules. An estimated 20% of the 660 Gt dissolved organic carbon in the ocean pool consists of structurally uncharacterized oligosaccharides, which appear to resist microbial degradation (Aluwihare et al., 1997). Current technologies lack the sensitivity and molecular resolution to identify these oligosaccharides. Here, we adapted graphitized carbon chromatography to extract and separate marine oligosaccharides for liquid chromatography high resolution mass spectrometry analysis. Using a newly-developed *de novo* annotation tool, we found 110 oligosaccharide structures in surface and deep ocean seawater at two distant locations in the North Atlantic Ocean. One group of the detected oligosaccharides was found only in surface seawater and consisted of larger and more abundant molecules detected by our analysis. A second group of smaller, less abundant oligosaccharides was detected in both the surface and deep ocean seawater of both sampled locations. The composition of oligosaccharides differed between the surface and deep ocean, with deep ocean samples relatively enriched in hard-to-metabolize deoxy-sugars, and xylose, amino sugars and uronic acids compared to simple hexoses. Notably the deoxy-sugar fucose constituted 35-40% of the monomers in deep-sea oligosaccharides, twice the percentage in surface ocean oligosaccharides. The ubiquity of deep ocean oligosaccharides indicates that they represent a preserved fraction of the carbohydrate pool. Their enrichment in specific monosaccharides suggests selective preservation of fucose-rich oligosaccharides in the deep ocean.

## Introduction

Photosynthetically fixed carbon dioxide enters the marine carbon cycle in the form of carbohydrates (1). Combined into polysaccharides, algal carbohydrates store energy, compose the cell wall, and form the extracellular mucilage layer (2). Heterotrophic bacteria use polysaccharides as a carbon and energy source, thereby remineralizing the stored carbon. The digestion of polysaccharides by heterotrophic bacteria generally proceeds via extracellular, enzymatic cleavage into oligosaccharides (3–5), which bacteria can then import into the cell using a diversity of transporters (6–9). Many of these transporter proteins are detected throughout the water column, suggesting effective turnover of algal carbohydrates in the ocean (10–13). However, quantitative measurements of bulk dissolved organic carbon (DOC) indicate that carbohydrate-like structures comprise 25-35% of the 660 Gt DOC in the ocean (14–17), with linkage analysis estimating the average carbohydrate length as 5 monomers (18). Moreover, carbohydrates can have average radiocarbon ages between one and three years in the surface ocean (19). Despite abundant heterotrophic bacteria with degradative capacity, carbohydrates - including oligosaccharides - persist throughout the water column (18, 20–23), indicating unexpected mechanisms that impede or prevent bacterial recycling. Determining factors that might contribute to this persistence requires higher structural resolution of oligosaccharides than is currently possible.

Recent introduction of biocatalytic and antibody-based methods for structure-specific analysis of polysaccharides revealed a diversity of structural motifs persisting in the ocean (24–27). Laminarin was found to account for approximately half of the organic carbon in diatom-containing particles (24, 28). Polysaccharides such as arabinogalactans, mannans and sulfated fucans were continuously present in dissolved organic matter during a diatom-dominated bloom in the North Sea (27). The sulfated fucans, which were shown to be secreted by diatoms, aggregated and accumulated in particulate organic matter, thereby providing a potential pathway for carbon sequestration (27, 29). Using antibodies and biocatalytic assays, the brown algae *Fucus vesiculosus* were also shown to secrete fucoidan at a rate of about 0.3% of their biomass per day (26). Antibody detection in sediments and slow degradation rates support considerable environmental stability of sulfated fucans (30–32). While biocatalytic and antibody-based methods have advanced our understanding of the role of carbohydrates in the marine carbon cycle, they are focused specifically on polysaccharides.

Methods are missing in particular for the extraction (**Table S1**) and analysis of smaller oligosaccharides, which likely represent the majority of dissolved carbohydrate carbon in the ocean (18). While more recalcitrant polysaccharides have been identified in the surface ocean (27), methodological obstacles obscure whether certain oligosaccharide structures have higher stability and, thus, a higher potential for carbon sequestration. Understanding of the cycling of specific marine oligosaccharides could be critical to, for example, accounting for dissolved organic carbon exported from macroalgae farms and forests, or from phytoplankton blooms (33–36). Structure-specific tracing of oligosaccharides (18) yielded from polysaccharides would represent a significant step forward.

To identify oligosaccharides in the dissolved organic matter pool, we developed methods that can isolate oligosaccharides from seawater for structural analysis. We started with a graphitized carbon chromatography solid phase extraction step (GCC-SPE) (37, 38), which allowed us to overcome low concentrations of oligosaccharides in the complex seawater salt matrix and to remove other molecules that could cross-react with oligosaccharides during sample processing (39–41). Using GCC-SPE, we processed surface and deep ocean samples obtained during two simultaneous research cruises in the North Atlantic Ocean. Liquid chromatography high resolution mass spectrometry analysis (LC-HRMS) with deep learning assisted denoising and *de novo* annotation revealed intact oligosaccharides across depths, which were confirmed by monosaccharide specific quantification. Oligosaccharide structures detected in deep ocean extracts were found at both sampling sites and, at higher concentrations, in the surface extracts. Compared to surface-specific oligosaccharides, deep ocean oligosaccharides were smaller, less abundant, and enriched in fucose and other unidentified deoxy sugars. Our results suggest the selective preservation of these small oligosaccharides, comprised of harder-to-metabolize monosaccharides, in the ocean.

## Results

Surface seawater from the deep chlorophyll maximum and deep ocean seawater was sampled south of Newfoundland (EN683) and north-west of Ireland (MSM107) in May/June 2022 (**Fig. 1A**). Oligosaccharide extractions were carried out on the cruises within the same week. Salinity and potential temperature measurements suggest that the two North Atlantic cruises sampled distinct deep ocean water masses (**Fig. 1C**). They characterize seawater sampled at 4,100 m depth off Newfoundland as North Atlantic Deep Water (NADW) influenced by Danish Straits Overflow Water (12, 42, 43) and seawater sampled at 2,300 m depth near Ireland as NADW influenced by Iceland-Scotland Overflow water (44). Each cruise collected 3 surface and 3 deep ocean seawater samples of 600 mL volume at one station, and 2 artificial seawater blanks, from which oligosaccharides were extracted using GCC-SPE. For oligosaccharide analysis by LC-HRMS, extracts were dried and dialyzed achieving a concentration factor of 400-fold relative to seawater.

**Figure 1.**
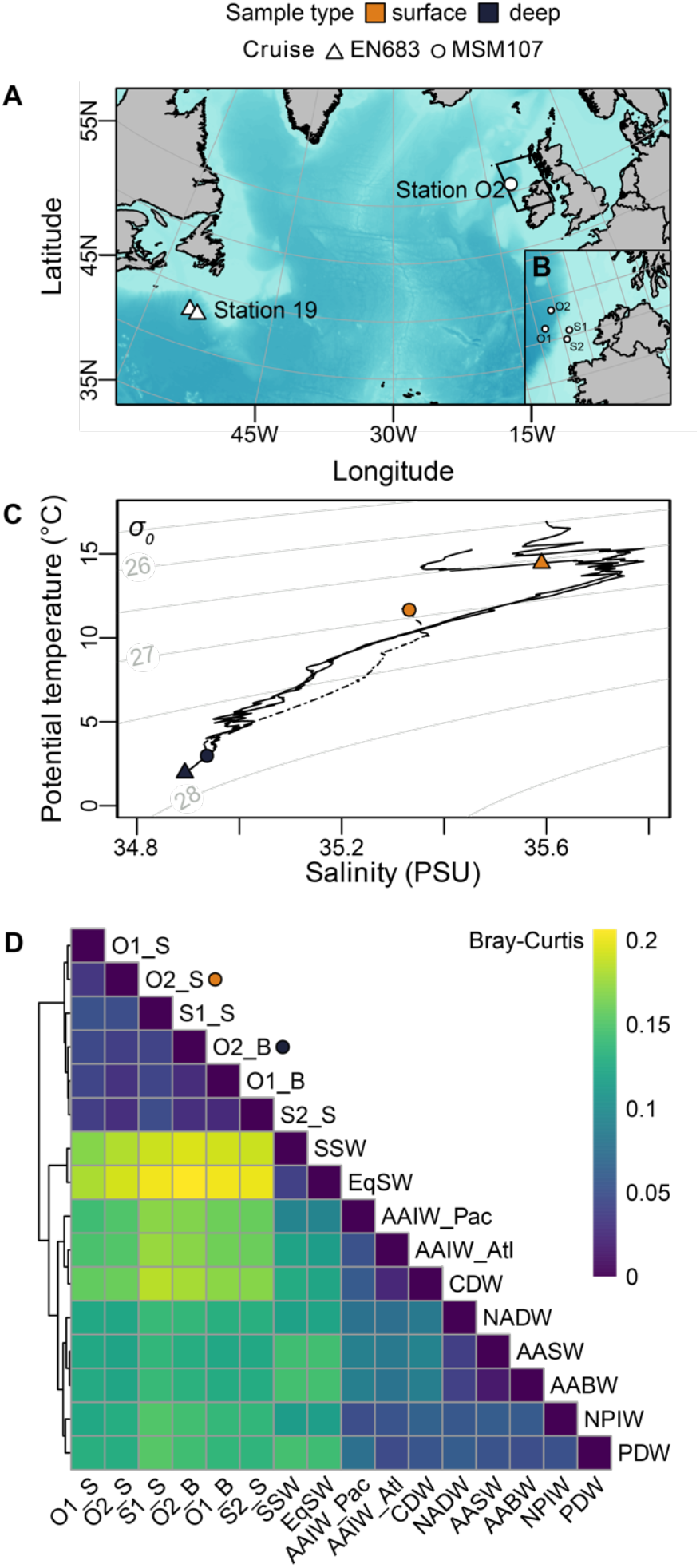
Distinct water masses sampled south of Newfoundland and northwest of Ireland. **A)** Sampling locations for oligosaccharides off the coast of Newfoundland during cruise EN683 (triangles) and northwest of Ireland during cruise MSM107 (circle). Bathymetry data is from NOAA (2022), and plotted with the R packages ‘marmap’ (50) and ‘oce’ (51). **B)** Locations of four stations sampled on MSM107 northwest of Ireland. Low molecular weight and hydrophobic dissolved organic matter was extracted by PPL solid phase extraction (SPE-DOM) from these seawater samples and analyzed by FT-ICR-MS. **C)** Physical parameters of the sampled water masses based on casts at station 19 and station O2. Colored points indicate depths at which seawater was sampled. **D)** Bray-Curtis dissimilarity matrix of 5,263 molecular formulae and corresponding FT-ICR-MS signal intensities of SPE-DOM from seawater collected on MSM107 and on previous expeditions (48). Dendrogram shows hierarchical clustering. O1, O2, S1 and S2 denote MSM107 stations (B), and ‘S’ and ‘B’ surface and bottom water. SSW = Subtropical Surface Water; EqSW = Equatorial Surface Water; AAIW_Pac = Antarctic Intermediate Water (Pacific); AAIW_Atl = Antarctic Intermediate Water (Atlantic); CDW = Circumpolar Deep Water; NADW = North Atlantic Deep Water; AASW = Antarctic Surface Water; AABW = Antarctic Bottom Water; North Pacific Intermediate Water; PDW = Pacific Deep Water.

Additionally, 4 surface and 2 deep samples were taken near Ireland to characterize the organic carbon environment of this sampling region **(Fig. 1B**). No such samples are available for the station near Newfoundland. Elevated dissolved organic carbon (DOC) concentrations at depth, typical for newly formed NADW, suggest recent deep water formation (45) (**Table S8**). Dissolved organic matter was extracted with PPL-SPE (SPE-DOM), which extracts up to 65% of the bulk, but retains neither large, more hydrophilic components such as carbohydrates nor small amino acids or organic acids including acetate (46, 47), and analyzed by FT-ICR-MS (**Fig. S7, Table S9**). Molecular formulae were assigned to 5,263 peaks, with 3,289 shared among all samples that represented 98.12% ± 0.48% (mean ± SD) of the total ion intensity of the samples. The samples collected here were molecularly more similar to each other than to samples from the global dataset, likely due to regional specificities and possible analytical variability between the two studies. Furthermore, the strong molecular similarity of our surface and deep samples is indicative for deep ventilation. Accordingly, our samples were molecularly more similar to other subpolar (>40 N and S) and polar (>50 N and S) latitude samples, compared to other oceanic water masses (48) (**Fig. 1D**). Elevated DOC concentrations and similar compositions of dissolved organic matter regardless of sampling depth indicated recent formation of the deep water sampled near Ireland (49).

### Oligosaccharide features occur in surface and deep ocean

The processed GCC-SPE extracts were analyzed using LC-HRMS, which can separate and detect intact oligosaccharides. Features in retention time and mass-to-charge ratio (*m/z*) space were computationally identified by pre-processing and deep learning-assisted denoising (52) (**Fig. 2A**). In a principal component analysis (PCA) of samples based on log2-transformed intensities of the 4,786 identified features, the first two components accounted for 43% of the total variation (**Fig. 2B**). The first dimension reflected sampling region, while the second separated surface, deep ocean, and processing blanks. Separation of sample groups in the PCA indicated successful extraction and detection of 1000s of dissolved seawater compounds, but did not specifically provide information about the presence of marine oligosaccharides.

**Figure 2.**
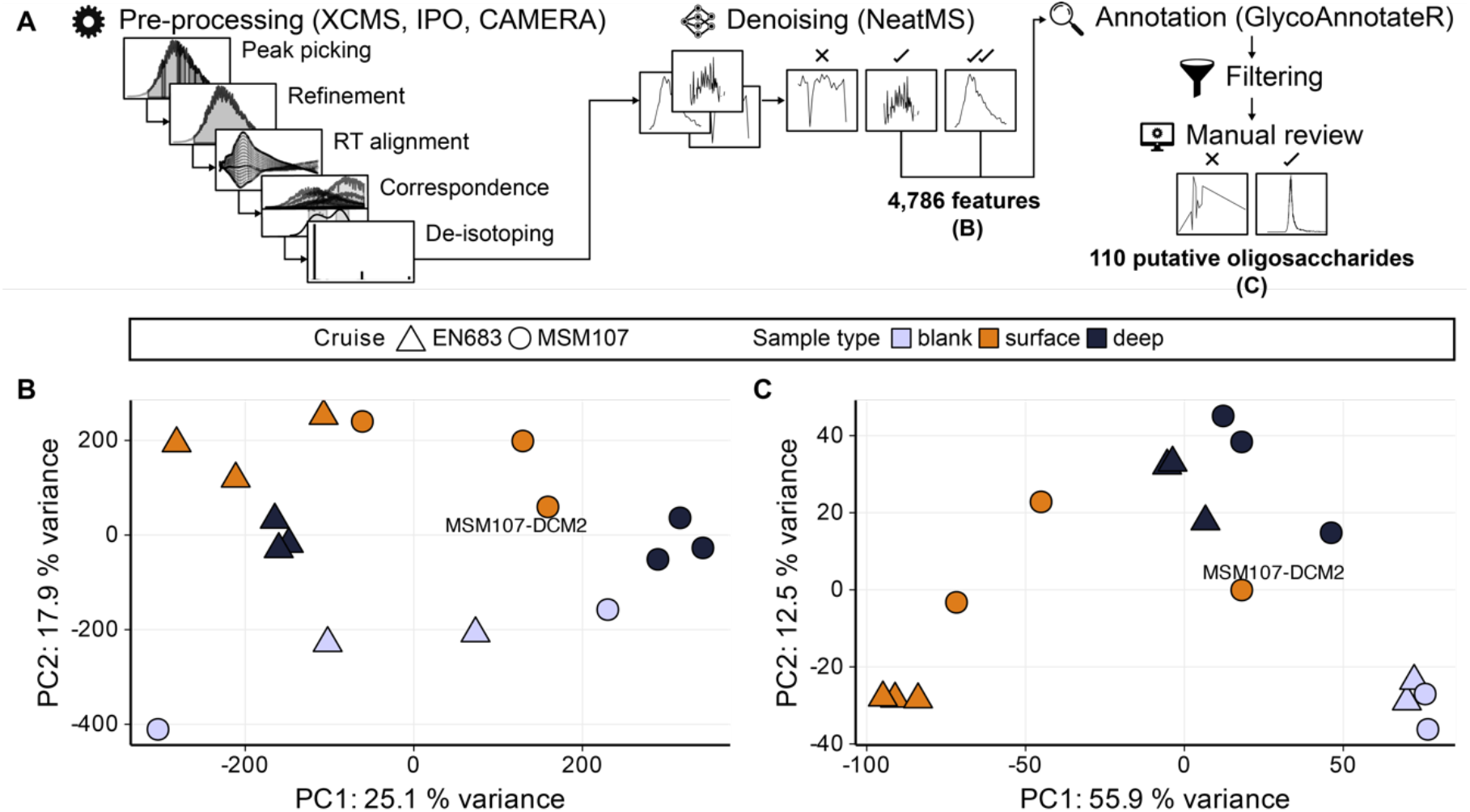
Putative oligosaccharides extracted from seawater reflect sampling depth. **A)** Liquid chromatography high resolution mass spectrometry data were pre-processed with XCMS (55) using IPO-optimized parameters (56) to generate features. ^13^C-isotopes identified with CAMERA (57) were removed. Identified MS1 peaks were then classified as noise (cross), low quality (one tick) and high quality (two ticks) by NeatMS (52). Analysis continued with low- and high-quality peaks, which corresponded to 4,786 features. These features were annotated (https://github.com/margotbligh/GlycoAnnotateR) to identify putative oligosaccharides. Annotated features were filtered to remove features abundant in the processing blanks (artificial seawater). Extracted ion chromatograms of the remaining features annotated as potential oligosaccharide ions were manually reviewed, with peaks across 110 ions designated as quality signals that represent putative oligosaccharides. **B)** Principal component analysis performed on log2-transformed integrated intensities of the 4,786 quality features. Light purple symbols correspond to processing blanks, orange symbols to surface extracts (deep chlorophyll maximum) and black symbols to deep ocean extracts, with shapes depicting the sampling expeditions. The labelled point MSM107-DCM2 is likely an outlier. **C)** Principal component analysis for the 110 features retained after annotation, filtering and manual quality classification.

Accordingly, we filtered our feature list to identify marine oligosaccharides, yielding 110 putative oligosaccharide features across sample groups. Features were annotated by comparison of observed and theoretical oligosaccharide ion *m/z* values derived from a combinatorial approach to ‘build’ chemically possible oligosaccharide structures (https://github.com/margotbligh/GlycoAnnotateR) (**Fig. 2A**). Of the 4,786 identified features, approximately 7% or 349 features were annotated as putative oligosaccharides. Our pipeline involved a successively narrowing stringency, with data pre-processing and denoising purposefully performed with parameters that maximized the number of signals retained, to include low intensity but true signals. Features were further filtered to remove those abundant in blanks, which controlled for contaminations from artificial seawater preparation, extraction columns and sample processing. This computational filtration step selected against any oligosaccharides contained in the sea salts used to recreate ionic strength in the artificial seawater blanks and returned 110 putative oligosaccharides. In a PCA of samples based on the log2-transformed intensities of the 110 putative oligosaccharides, principal components 1 and 2 explained 68.5% of variation (**Fig. 2C**). One surface sample from MSM107 (MSM107-DCM2) likely represented an outlier. Seawater samples grouped by depth rather than region, indicating a pronounced effect of water mass age.

The identified 110 putative oligosaccharides could each comprise a single chemical structure or multiple isomers (**Fig. S8, S9**). Incomplete resolution of isomers is inherent to carbohydrate LC-MS due to the high potential for stereochemical diversity (53,54). Despite GCC-SPE, oligosaccharide concentrations were too low to achieve additional characterization with fragmentation spectra generated via tandem mass spectrometry. In addition, the number of hypothetical annotations per feature corresponding to possible compositions for a detected *m/z* ranged between 1 and 58 with an average of 9.5 ± 12.9. Nonetheless, the LC-HRMS analysis showed that putative oligosaccharides could be separated and traced across samples. This yielded for each GCC-SPE extract relative abundances of putative oligosaccharides based on integrated peak areas.

### Small oligosaccharides detected across depths

Clustered by their abundance across 16 samples, the 110 putative oligosaccharides separated into four groups with distinct size distributions. Multiple methods supported the use of *k=4* in *k*-means clustering based on log2-transformed intensities across samples (**Fig. S10**). The first dimension of a PCA performed on putative oligosaccharides accounted for 45.8% of variation and separated clusters 1 and 2 from clusters 3 and 4 (**Fig. 3A, B**). Dimensions 2 and 3 separated clusters 1 and 2, and clusters 3 and 4 respectively (**Fig. 3A, B**). In our analysis, *m/z* is a proxy for size, as detected oligosaccharide ions likely carried only a single charge (*z* = 1) (see Supplementary Methods). Putative oligosaccharides in clusters 3 and 4 were significantly smaller than oligosaccharides in cluster 1 (**Fig. 3C**; pairwise Wilcox signed-rank tests, Benjamini-Hochberg adjusted *p*-values < 0.005). The distinct size distributions of putative oligosaccharide clusters suggested that these groupings may be related to physical or biological features of the ocean.

**Figure 3.**
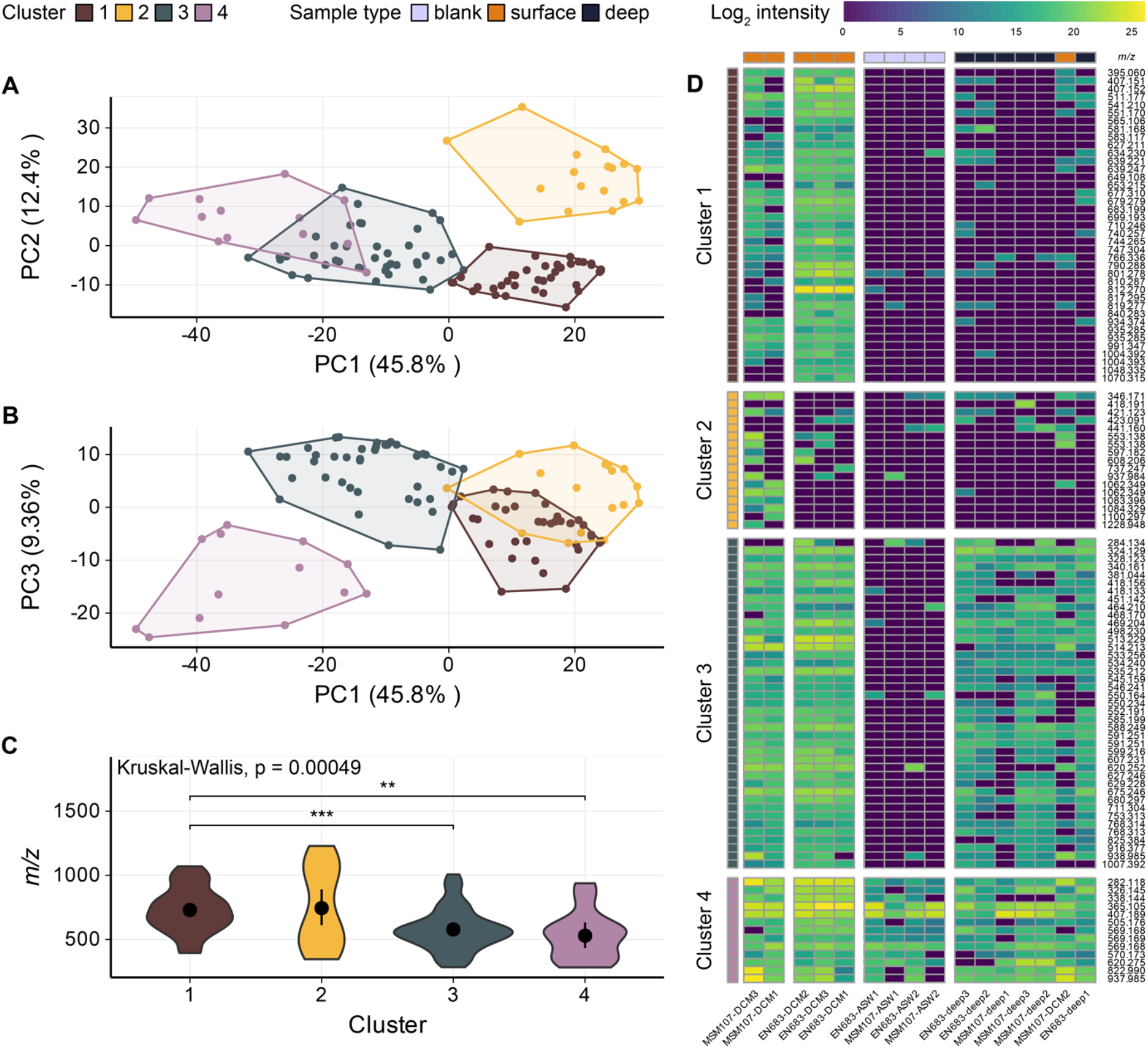
Putative oligosaccharides are either larger and limited to surface samples, or smaller and ubiquitous. Putative oligosaccharides and samples were grouped based on log2-transformed intensities by *k*-means clustering (*k* = 4 for clustering of both features and samples, **Fig. S10, S11**). **A)** Separation of putative oligosaccharides along the first two components of a principal components analysis explained ∼63% of the variation. **B)** Visualization of principal components 1 and 3 of the same analysis shows separation of clusters 3 and 4. **C)** The clusters have significantly different *m/z* distributions (p < 0.001). Clusters 3 and 4 have significantly (adjusted p < 0.005) lower *m/z* distributions than cluster 1. Black points represent means of clusters, and lines represent 95% confidence intervals of the means from nonparametric bootstrapping. **D)** A heatmap of the log2-transformed feature intensities shows the association of clusters with sample groups. Both rows and columns are ordered by *k*-means cluster identities with gaps between clusters. Clusters 1 and 2 are surface-specific. Cluster 3 features are present in both surface and deep samples. Cluster 4 features appear in all samples including processing blanks (artificial seawater) and likely represent contaminants. The *m/z* of each feature is shown on the right.

Mapping out clusters sample-by-sample and performing *k*-means clustering of samples (**Fig. S11**) revealed that, while larger putative oligosaccharides were specific to surface seawater extracts, smaller putative oligosaccharides recurred across sampling locations and depths (**Fig. 3D**). The 41 putative oligosaccharides in cluster 1 had an average *m/z* of 729.401 ± 176.090 (SD) and occurred in surface waters from both locations, but only sporadically in deep ocean samples. The 17 putative oligosaccharides in cluster 2, which primarily occurred in surface samples collected near Ireland, distinguished the sampling locations (**Fig. 3D**). Cluster 3 harbored 39 putative oligosaccharides with an average *m/z* of 578.127 ± 164.106 and occurred consistently in surface and deep ocean extracts, with intensities on average ∼50% higher in surface samples. The 13 putative oligosaccharides in cluster 4 were present in all samples including blanks. Saturation curves support the notion that, with the detection limits intrinsic to the analysis, surface samples contain more oligosaccharides than deep ocean samples (**Fig. S12**). Thus, larger oligosaccharides were unique to surface extracts, while smaller oligosaccharides exist in both surface and deep ocean seawater, albeit at lower concentrations in the latter.

### Relative monosaccharide abundances in oligosaccharides differ between depths

We used an established approach of quantifying monosaccharides (**Fig. S6, Table S7**) after acid hydrolysis to verify oligosaccharide extraction with GCC-SPE and compare their compositions (**Fig. 4**) (58, 59). Due to a glucose contamination in processing blanks, likely from the sea salts used to make the artificial seawater, glucose could not be confidently quantified in seawater extracts, but other sugars were unaffected (**Fig. 4A**). After excluding glucose and subtracting processing blank values for all monosaccharides, monosaccharides in surface oligosaccharides summed to 0.17-0.26 µM carbon compared to 0.04-0.08 µM carbon in deep ocean oligosaccharides. Fucose, galactose, xylose, ribose, mannose, rhamnose and arabinose were the most abundant monomers detected, with fucose consistently constituting at least 20% after blank subtraction (**Fig. 4B**). In addition to the identified monosaccharides, three deoxyhexoses and five uronic acids with retention times distinct from standards occurred in extracted oligosaccharides (**Fig. 4D, E, Fig. S13**). Using monosaccharide quantification as an independent method supported LC-HRMS results by demonstrating convergence of monosaccharide composition between sampling locations and confirming differences in absolute concentrations between depths.

**Figure 4.**
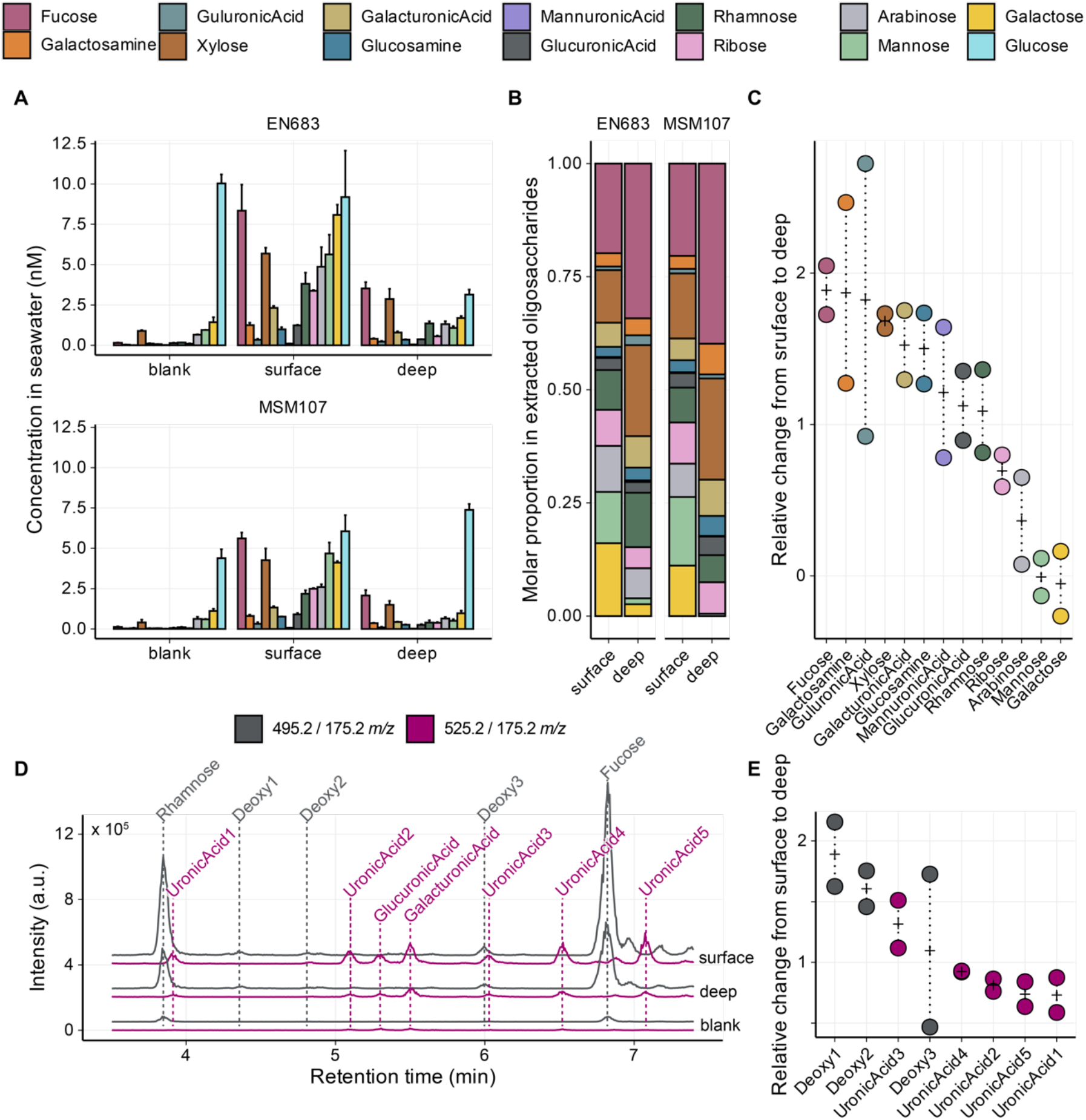
Extracted oligosaccharides indicate enrichment of fucose- and other deoxyhexose-containing oligosaccharides in the deep ocean. **A)** Monosaccharide concentrations (nM) in processing blanks (artificial seawater) and extracts from surface (deep chlorophyll maximum) and deep ocean seawater from south of Newfoundland (EN683) and northwest of Ireland (MSM107). Bars correspond to the mean of three replicates and error bars to the standard error. Glucose contamination of processing blanks likely came from the sea salts used to make the artificial seawater. Processing blank values were subtracted for B and C. **B)** Mean molar contribution of monosaccharides for each sampling site and depth. **C)** Change in molar proportion of monosaccharides detected in deep ocean extracts compared to surface extracts for each sampling site. Crosses indicate the means across the sites. **D)** Example chromatograms of precursor/product ion pairs of deoxyhexoses (Deoxy) and uronic acids (UronicAcid). Labels with numbers indicate unidentified monosaccharides. **E)** Relative changes in response ratios of unidentified monosaccharides detected in deep ocean extracts compared to surface extracts for each sampling site. These unidentified monosaccharides could not be quantified without analytical standards and their relative contributions are only in comparison to each other.

The monosaccharide composition of the extracted oligosaccharides differed between depths. Here, relative abundance refers to the molar proportion of a monosaccharide after subtraction of processing blank values (**Fig. 4B**). The relative abundance of fucose, galactosamine, xylose, galacturonic acid and glucosamine in deep ocean oligosaccharides exceeded their relative abundance in surface oligosaccharides at both sites (**Fig. 4C**). Guluronic acid, mannuronic acid, glucuronic acid and rhamnose varied between relative enrichment or depletion at depth depending on the sampling site. Ribose, arabinose, mannose and galactose were relatively less abundant at depth compared to surface for both sites. Mannose and galactose concentrations were higher in our processing blanks than deep ocean samples, resulting in apparent fold-changes in relative abundance below 0. Two unidentified deoxyhexoses showed increases in relative abundance similar to fucose (**Fig. 4E**), although these values are only relative to the response ratios of the other unidentified deoxyhexoses and uronic acids. Fucose not only showed the greatest increase in relative abundance by depth (1.89-fold ± 0.23) of all monosaccharides, but also had the highest average relative abundance at both depths (20.11% ± 0.41% and 38.00% ± 5.35% for surface and deep respectively).

## Discussion

This study detected oligosaccharides in surface and deep ocean seawater from two sites in the North Atlantic Ocean, an analysis enabled by integrating graphitized carbon extraction (GCC-SPE) with liquid chromatography high resolution mass spectrometry (LC-HRMS). This analysis revealed two groups of putative oligosaccharides in sampled seawater extracts: larger putative oligosaccharides in surface ocean extracts, only sporadically present in deep ocean extracts, and smaller putative oligosaccharides ubiquitous detected across sites and depths (**Fig. 3**). Approximately one third of surface oligosaccharide concentrations were detected in deep ocean seawater (**Fig. 3, 4**). Monosaccharide compositions differed between surface and deep ocean oligosaccharide extracts, with structurally more complex monosaccharides relatively enriched in deep ocean samples. Our findings reveal the presence of small oligosaccharides throughout the water column in the North Atlantic Ocean and suggest the selective preservation of oligosaccharides comprised of hard-to-metabolize monosaccharides.

Purification and chromatographic separation enabled the discrimination of oligosaccharides that persist in the ocean on timescales longer than seasonally. Chromatographic separation removes the potential for cross-reactions that may occur during bulk analyses or ultrafiltration (39–41), in which all molecules above a size threshold are concentrated. Using chromatography for isolation and separation of marine dissolved carbohydrates allowed us to confidently discriminate between distinct molecules. Complementary, molecular SPE-DOM analyses on samples from the Irish Sea revealed only minor differences between the composition of surface and bottom water extracts and suggest recent deep water formation (**Fig. 1D**). Rapid cycling of seasonally produced dissolved organic molecules in the SPE-DOM molecular window has been proposed for high latitude marine systems (49). In contrast to this, we observed that larger, more abundant oligosaccharides were surface specific (**Fig. 3**), likely representing an imprint of spring productivity in the surface ocean. The ubiquity of smaller oligosaccharides in surface and deep ocean samples suggests that these compounds are, unlike labile low molecular weight compounds, cycled on timescales at least longer than seasonally.

These smaller, less abundant oligosaccharides identified in the deep ocean extracts likely represent persistent remnants of surface productivity. Transport of surface-produced oligosaccharides to depth within subducted or downwelled water masses would imply 2 to 100 years since surface contact (60-62). Alternatively, oligosaccharides could be transported by particles sinking out of the surface ocean at 0.1-200 m d^-1^ taking days to centuries to reach the deep ocean (63-65). Finally, oligosaccharides could stem from *in situ* production by microbial communities in surface and deep ocean water, with higher concentrations at surface due to higher microbial activity. *In situ* production of identical oligosaccharides at contrasting depths and two locations ∼3,000 km apart appears unlikely in such different environmental conditions with distinct microbial communities (12, 66-68). Instead, the absence of any oligosaccharides unique to deep ocean samples, lower concentrations of oligosaccharides in the deep ocean and their recurrence across locations suggest that deep ocean oligosaccharides are surface-produced molecules with slow turnover rates (**Fig. 2C, 3C, 4)**.

We recognize that extending this molecular picture to wider regions of the world’s oceans will require further sampling, for example, along a more highly-resolved depth gradient, as well as sampling in other regions. Improvement of the sample blank procedures to avoid contaminations by commercial sea salts will like improve data by permitting inclusion of additional hexose oligosaccharides and glucose quantification in future studies. We also note that the incomplete separation of isomers (**Fig. S8**) means that reported feature numbers only constitute a lower boundary of marine oligosaccharide diversity. Establishing an upper boundary may require other methods that enable separation and characterization of isomers, for example ion mobility (54, 69).

Nonetheless, monosaccharide compositions determined here compare well with previous studies of marine carbohydrates. Up to 20% of total dissolved organic carbon consists of oligosaccharides with an average length of five monomers based on linkage analysis of ultra-filtered dissolved organic matter (>1 kDa, UDOM) (18). Our oligosaccharide extracts matched the monosaccharide compositions reported for UDOM (**Fig. 4A, B**) (18, 20). One difference, however, is that we found ribose to constitute 7.8 ± 2.8% of total quantified monosaccharides, while earlier studies of marine carbohydrates reported ribose below their detection limit (20). Stronger acid conditions in previous studies may have promoted a higher conversion rate of ribose to furfural (70). Our detection of ribose, which is about tenfold more reactive than other neutral monosaccharides (71), may alternatively indicate chromatographic separation before analysis reduces cross-reactions during sample processing. The observation that our monosaccharide compositions align with previous reports of marine carbohydrates support extrapolation from patterns observed here to the dissolved carbohydrate pool.

Our results demonstrate relative enrichment with depth of deoxyhexoses over hexoses and to a lesser degree of uronic acids over hexoses, likely reflecting differences in ease of metabolism of these monomers. This increase in molar proportion from surface to depth of fucose and unidentified deoxyhexoses (**Fig. 4C, D**) matches previous observations of shifts in monosaccharide composition of bulk carbohydrates with depth (18, 19, 72/74) (**Table S10**) and during diagenesis (75, 76). Hexoses connect well into constitutive cellular pathways facilitating their catabolism as exemplified by the observation that the hexoses mannose and galactose exhibited approximately 95% to 100% depletion at depth (**Fig. 4**). Uronic acids, which are depolymerized in cascades involving polysaccharide lyases and enter catabolism after several enzymatic steps, showed moderate depletion (**Fig. 4C, E**) (5, 77, 78). Deoxyhexoses including fucose and rhamnose require dedicated catabolic pathways only found in few bacteria (79. 80). Compared to surface samples, molar proportions of fucose and two unidentified deoxyhexoses approximately doubled at depth, therefore comprising the most persistent monosaccharides (**Fig. 4C, E**). This greater persistence of oligosaccharides consisting of more complex and thus harder-to-degrade monosaccharides suggests their selective preservation.

Our techniques to concentrate dilute oligosaccharides from seawater and to distinguish structures could be applied in future studies to trace specific oligosaccharide structures in the ocean. While the sample volume used in this study (600 mL) represents a more than hundred-fold increase over the typical GCC-SPE volume of 0.5-3 mL (37, 38), the extraction volume is scalable due to the selective retention of target compounds and could yield sufficient oligosaccharide concentrations for tandem mass spectrometry (MS/MS) experiments. Initial identification and MS/MS characterization of target oligosaccharides, for example oligosaccharides yielded by hydrolysis of sulfated fucans (79, 81), would then allow for targeted tracing of those oligosaccharides in the ocean. Such tracing could enable species-specific accounting for DOC, for example during and after a phytoplankton bloom or in transects away from a macroalgae forest or farm. Despite their abundance in the DOC pool (18), oligosaccharides were not included in recent molecular-level studies of dissolved organic matter (86), primarily due to analytical restraints. Here, we developed techniques to overcome these restraints and found evidence for selective preservation of fucose-rich oligosaccharides in the North Atlantic Ocean.

## Conclusion

Here, we adapted a graphitized carbon solid-phase extraction method to capture short-chain carbohydrates - oligosaccharides - from seawater. The separation and identification of extracted oligosaccharides with LC-HRMS revealed that one group of larger oligosaccharides appears to be produced in the surface ocean, and is not present at depth, suggesting their degradation. A second group of smaller oligosaccharides was detected in all seawater samples including deep ocean samples, but yielded more intense signals at the surface. This pattern of occurrence indicates their production at the surface and persistence to reach the deep ocean. Persistent deep ocean oligosaccharides contained relatively more deoxyhexoses including fucose in comparison to surface oligosaccharides, which aligns with previous observations on bulk marine carbohydrates and suggests that preservation is selective. In summary, our findings suggest greater recalcitrance of specific carbohydrates, which are harder to metabolize, to allow a ubiquitous pool of recognizable oligosaccharides to persist in the ocean. The methods developed here could allow tracing of specific oligosaccharides in the dissolved organic carbon pool in future studies.

## Materials and Methods

### Sample acquisition

Surface samples originated from the deep chlorophyll maximum (called ‘surface’ in this study) and deep ocean samples from 4,100 and 2,300 m depth, respectively, from sampling locations at 43°N, 54°W south of Newfoundland and 55°N, 10W northwest of Ireland (**Table S2**). Physical parameters were recorded using a Seabird 911+ CTD profiler. Data were processed and binned with SBE Data Processing software (v7.26.7). Derived parameters were calculated using Ocean Data View (ODV; version 5.6.2) and the R package ‘oce’ v1.8-3 (51). Seawater was collected using 30 L Niskin (off Newfoundland) and 12 L Niskin (off Ireland) bottles attached to a rosette, transferred into acid-washed carboys and pre-filtered at 0.22 µm (Millipore Express PLUS Membrane Filter, hydrophilic polyethersulfone). Artificial seawater prepared from sea salts (Sigma, NutriSelect® Basic, lot SLBX3050) served as processing blanks.

### Oligosaccharide extraction

Oligosaccharides were extracted immediately after sampling using non-porous graphitized carbon Supelclean™ ENVI-Carb™ SPE cartridges (GCC-SPE, 60 mL, 10 g) following an adapted protocol for carbohydrate purification (**Fig. S1**) (83). The column volume (CV) was 60 mL. Cartridges were conditioned with 1 CV acetonitrile followed by 1 CV MilliQ-water. Samples of 600 mL filtered seawater or processing blanks were applied using gravity flow, facilitated by extending the reservoir using 3D printed cones. After four MilliQ-water washes of ½ CV each, oligosaccharides were eluted with 3x ¼ CV elution buffer consisting of 49.95: 49.95: 0.1 (v/v) MilliQ-water: acetonitrile: formic acid. Samples were partially dried by vacuum centrifugation then freeze-dried.

### Oligosaccharide analysis

To remove any residual salts prior to liquid chromatography high resolution mass spectrometry (LC-HRMS) analysis of intact oligosaccharides, extracts were dialyzed against MilliQ-water through 100-500 Da membranes (SpectraPor® Biotech), freeze-dried and resuspended in MilliQ-water. A mixture of aliquots from all samples served as quality controls. For LC-HRMS using a hydrophilic interaction liquid chromatography (HILIC) stationary phase, acetonitrile was added to samples to comprise 70% of the final volume, leading to a final concentration factor of 400-fold relative to seawater. In randomized order (**Table S3**), samples and standards (**Table S4**) were analyzed, with pooled quality controls (**Fig. S2**) every 5 injections, on an Accucore-150-amide HILIC column (Thermo Scientific, USA) coupled to a Thermo Fisher Q-Exactive Plus mass spectrometer in positive ionization mode.

### Identification of oligosaccharides

The obtained LC-HRMS spectra were centroided and converted to mzML format with msConvert (ProteoWizard), and pre-processed using the ‘XCMS’ (v4.0.2) package in R with IPO (v1.20.0) optimized parameters (**Table S5, Fig. 2A**) (55, 56, 84). Features annotated as ^13^C-isotopes by CAMERA (v1.56.0) after peak grouping by retention time were removed (57). Peaks were denoised by NeatMS (52). The classifier of the default NeatMS model was trained by transfer learning with a set of 2,000 manually classified and reviewed peaks (**Table S6, Fig. S3, Fig. S4**). Quality features were annotated as putative oligosaccharides by comparison with theoretical *m/z* values of de novo calculated compositions (https://github.com/margotbligh/GlycoAnnotateR), allowing carboxylic acid, deoxy, N-acetyl, O-acetyl, O-methyl, amino, anhydro-bridge, alditol and sulfate modifications for pentoses and hexoses with degrees of polymerization (DP) 2-6, and a mass deviation of 3 ppm. Features with integrated peak intensities greater than 1E8 (**Fig. S5**) or >60% of the maximum integrated intensity in any of the processing blanks were removed (85). Due to the relatively high false positive rate (0.09) at the model threshold (**Table S6**), remaining peaks were manually re-classified as quality or noise, returning 110 quality putative oligosaccharides.

### Monosaccharide quantification

Reconstituted extracts were transferred to pre-combusted (450°C, 4.5h) glass ampoules for hydrolysis in 1 M hydrochloric acid at 100°C for 24 h. Hydrolysates were dried by vacuum centrifugation to evaporate the acid, reconstituted in MilliQ-water and pH adjusted to >7 with 0.1 M sodium hydroxide. Standards consisted of a serially diluted mixture of monosaccharides (**Table S7**). Acid hydrolysates and standards were spiked with ^13^C-labelled glucose, galactose and mannose, and subsequently derivatized with 1-phenyl-3-methyl-5-pyrazolone (PMP) following (58). PMP-derivates were measured on a SCIEX qTRAP5500 and an Agilent 1290 Infinity II LC system equipped with a Waters CORTECS UPLC C18 column by multiple reaction monitoring (MRM) (59). Signal intensities were normalized to internal standards and calibrated against standard curves (**Fig. S6, Table S7**).

### Molecular characterization of SPE-DOM

Filtered seawater samples collected at four stations northwest of Ireland (**Table S2, Fig. 1B**) were acidified to pH 2 and hydrophobic, low-molecular weight dissolved organic matter, which does not include oligosaccharides, was solid phase-extracted using styrene divinylbenzene polymer cartridges (PPL) (46). This SPE-DOM was analyzed by Fourier transform ion cyclotron resonance mass spectrometry (FT-ICR-MS) on a Bruker Solarix XR (Bruker Daltonik GmbH, Bremen, Germany), equipped with a 15 Tesla superconducting magnet (Bruker Biospin, Wissembourg, France) and an electrospray ionization source (Apollo II ion source, Bruker Daltonik GmbH) in negative ionization mode at 4.5 kV (86). FT-ICR-MS spectra were calibrated and processed (see Supplementary Information for details), with molecular formulae assigned to peaks using the ICBM-Ocean processing pipeline (87). Processed spectra were averaged across technical (injection) duplicates and extraction triplicates, and compared to similarly processed FT-ICR-MS spectra from different water masses that are publicly available (48).

Detailed descriptions of the GCC-SPE protocol, NeatMS model training, LC-HRMS settings and procedures and data analysis, monosaccharide quantification, and SPE-DOM analysis can be found in the supplementary information.

## Acknowledgments

The authors thank the crews of R/V Endeavor during cruise EN683 and F/S Maria S. Merian during cruise MSM107, as well as technicians and colleagues at the Max-Planck-Institute for Marine Microbiology, including Alek Bolte, Katia Föll, Marvin Weinholz, Aman Akeerath Mundanatt, Nguyen Nguygen, Christopher Fitzgerald, Katrin Klaproth, Ina Ulber, Matthias Friebe and at the MARUM, especially Lennart Stock, and members of the mechanical workshop, namely Fabian Schramm. HBW, MBL, MLI, TD and JHH acknowledge support from the Cluster of Excellence initiative (EXC-2077–390741603), the DFG (HE 7217/1-1) and Heisenberg grant Glyco-Carbon Cycling in the Ocean), European Research Council grant C-Quest (47010790), the Simons Collaboration on Principles of Microbial Ecosystems collaboration (40200753) and the Max-Planck-Society. CA was supported by NSF OCE-2022952.

