## Supplementary for "Selective preservation of fucose-rich oligosaccharides in the North Atlantic Ocean"

### Supplementary figures

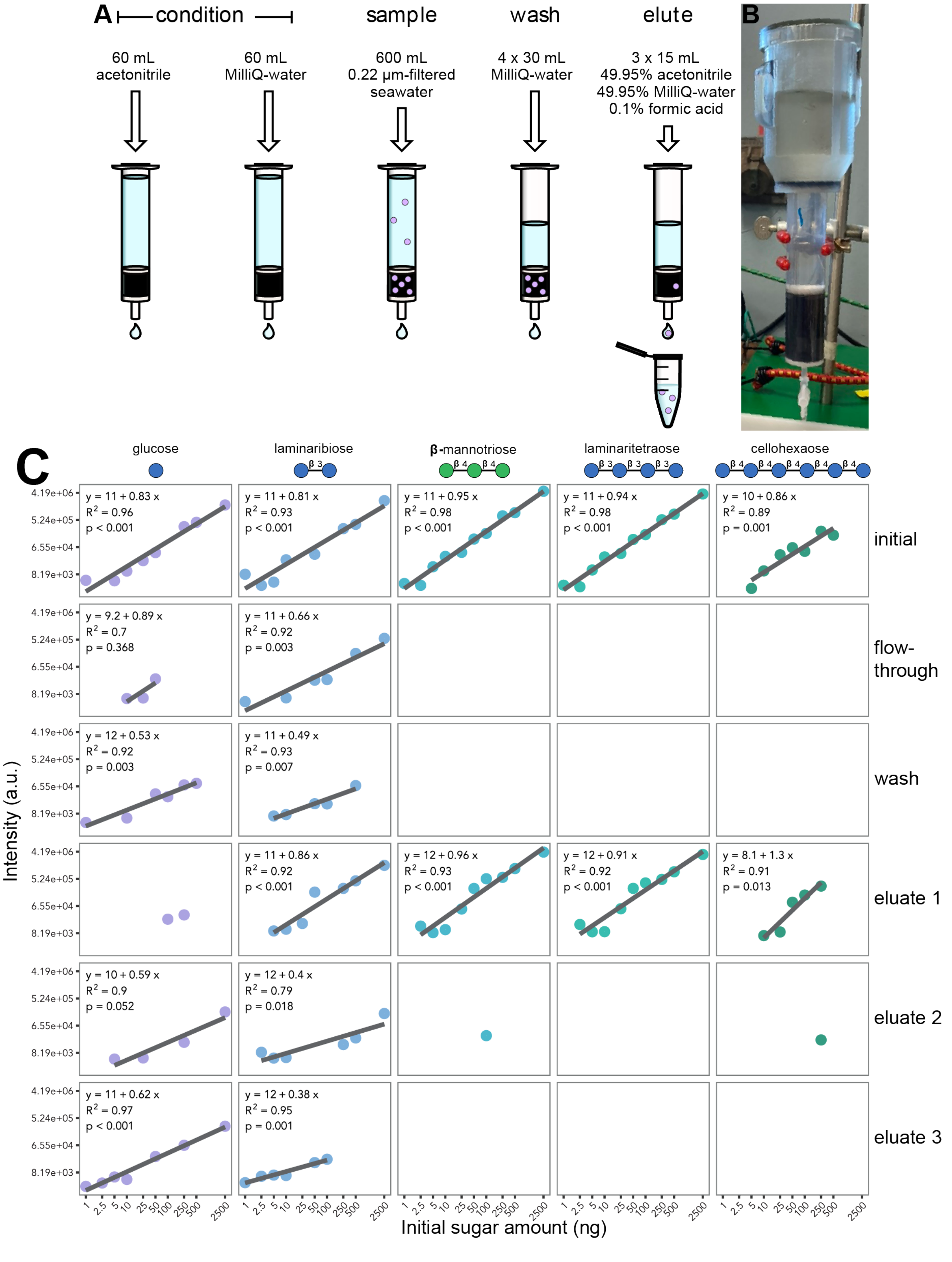

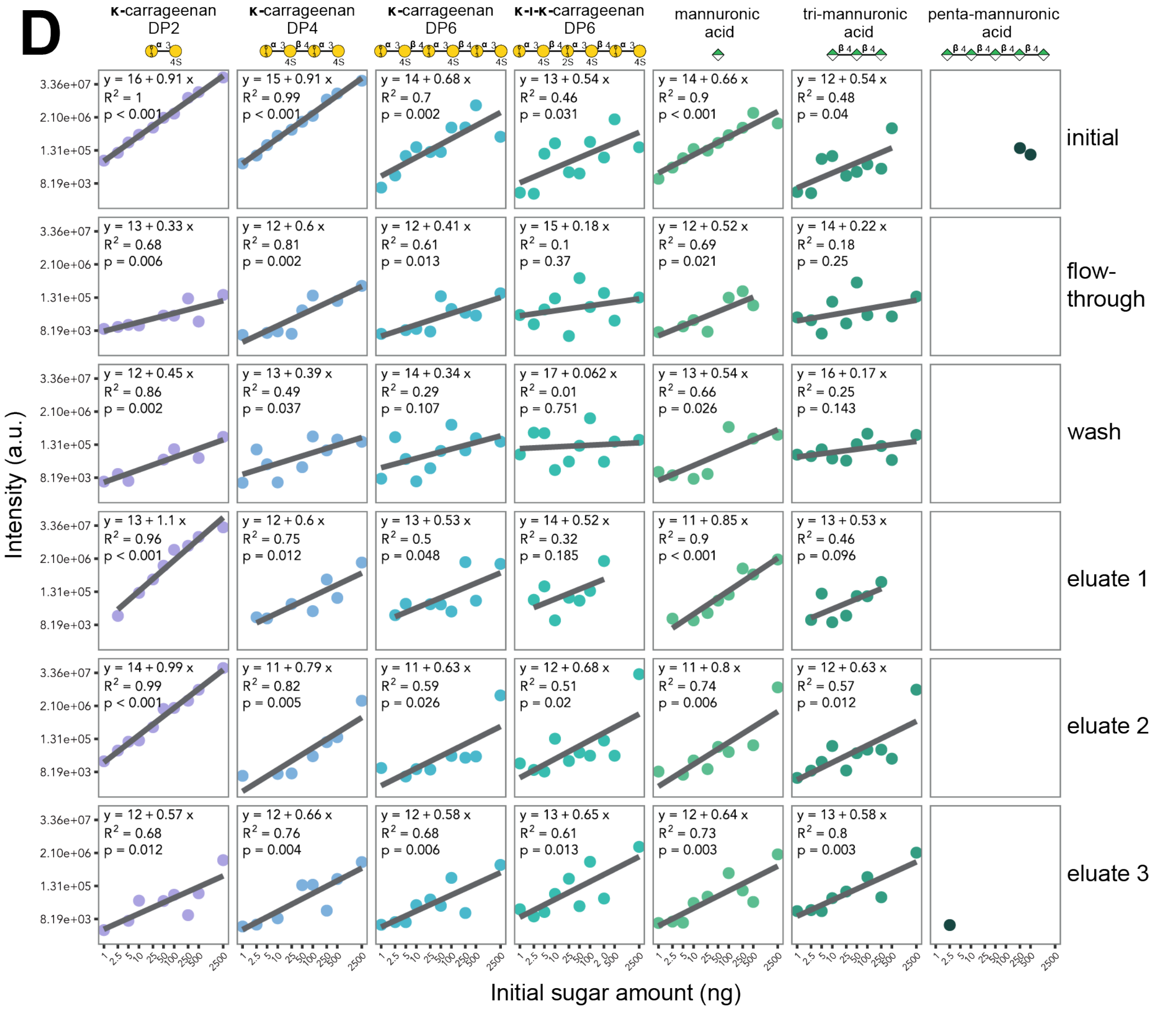


**Figure S1 | Non-porous graphitized carbon cartridge volumes extended with 3D-printed reservoirs. (A)** Protocol for extraction of carbohydrates from filtered seawater with graphitized carbon as a solid phase adapted from (1). **(B)** Cartridge volume was extended from 60 mL to 300 mL total. Reservoirs were soaked in acetonitrile (LC-MS grade) overnight before first use and MilliQ-washed five times in between uses. **(C-D)** Extraction of 1 - 2500 ng of neutral (C) and anionic (D) carbohydrates. Different amounts of commercial standards were dissolved in 3 mL MilliQ-water and extracted with 3 mL graphitized carbon cartridges. Initial, flow through, wash and three eluate fractions were collected and analyzed by direct infusion mass spectrometry on a Thermo Fisher Q-Exactive Plus. Spectra were acquired for 2 min while samples were infused at 5 µL/min and averaged. Both x (initial sugar amount) and y (signal intensity) axes are shown with log2 scales.

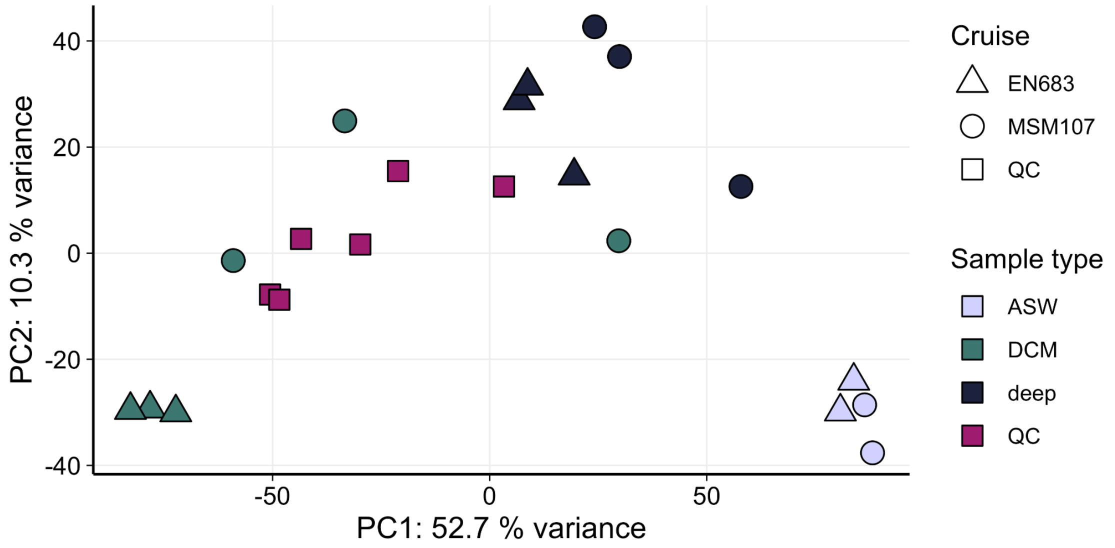


**Figure S2 | Pooled quality controls (QCs) are representative of seawater extracts based on putative oligosaccharides.** Principal component analysis (PCA) of the log2-transformed intensities of the 110 features identified as putative oligosaccharides. QCs were made by combining aliquots of all samples (seawater and artificial seawater extracts) and run every 5 injections.

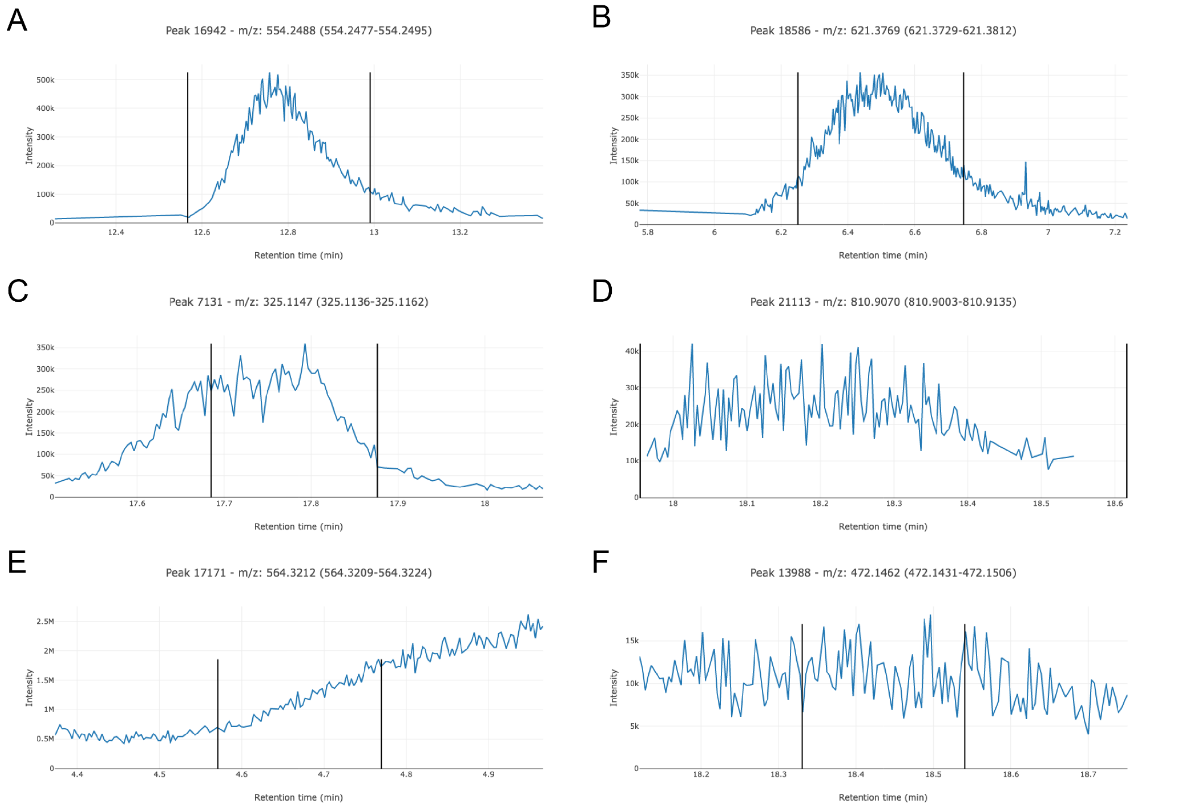


**Figure S3 | A set of 2000 peaks were manually classified as high or low quality or noise for training of the NeatMS model.** Example peaks (extract ion chromatograms) classified as high quality (A, B), low quality (C, D) and noise (E, F). *m/z* ranges: A = 554.2477-554.2495, B = 621.3729-621.3812, C = 325.1136-325.1162, D = 810.9003-810.9135, E = 564.3209-564.3224, F = 472.1431-472.1506.

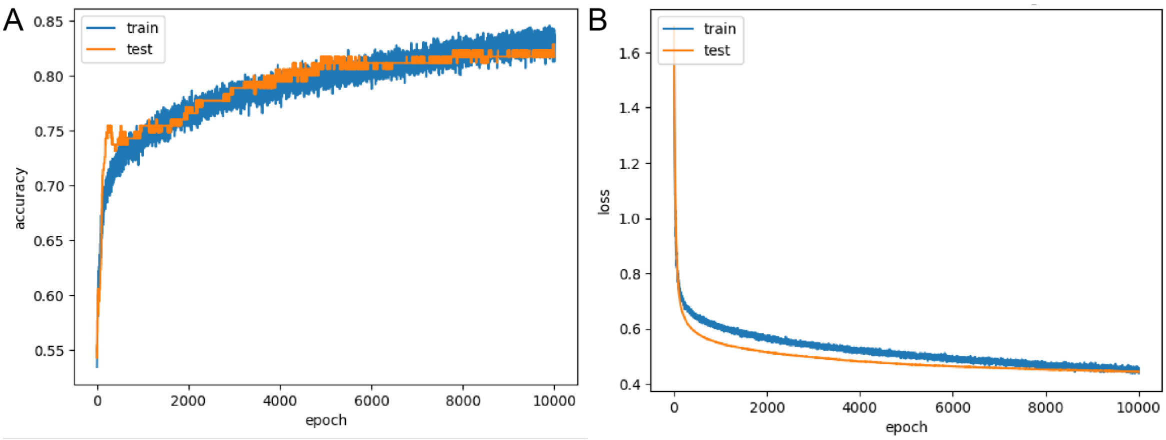


**Figure S4 | The classifier of the default NeatMS model was trained via transfer learning for 10,000 epochs at a low learning rate.** The learning rate was set to 0.000001 with “Adam” as optimizer. Plots show the accuracy (A) and loss (B) over 10,000 epochs for the training (blue lines) and test (orange lines) datasets.

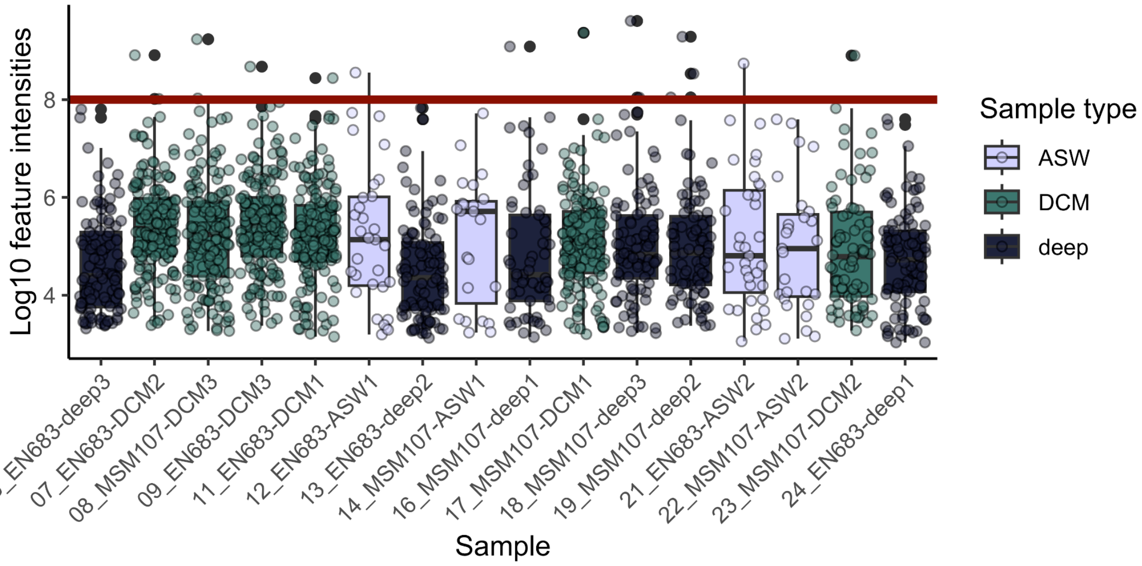


**Figure S5 | Features with integrated intensities greater than 1E8 were removed as potential outliers.** Integrated intensities (log10 scale) of individual features identified by pre-processing and after NeatMS denoising for each sample. The red line indicates the cut-off of 1E8.

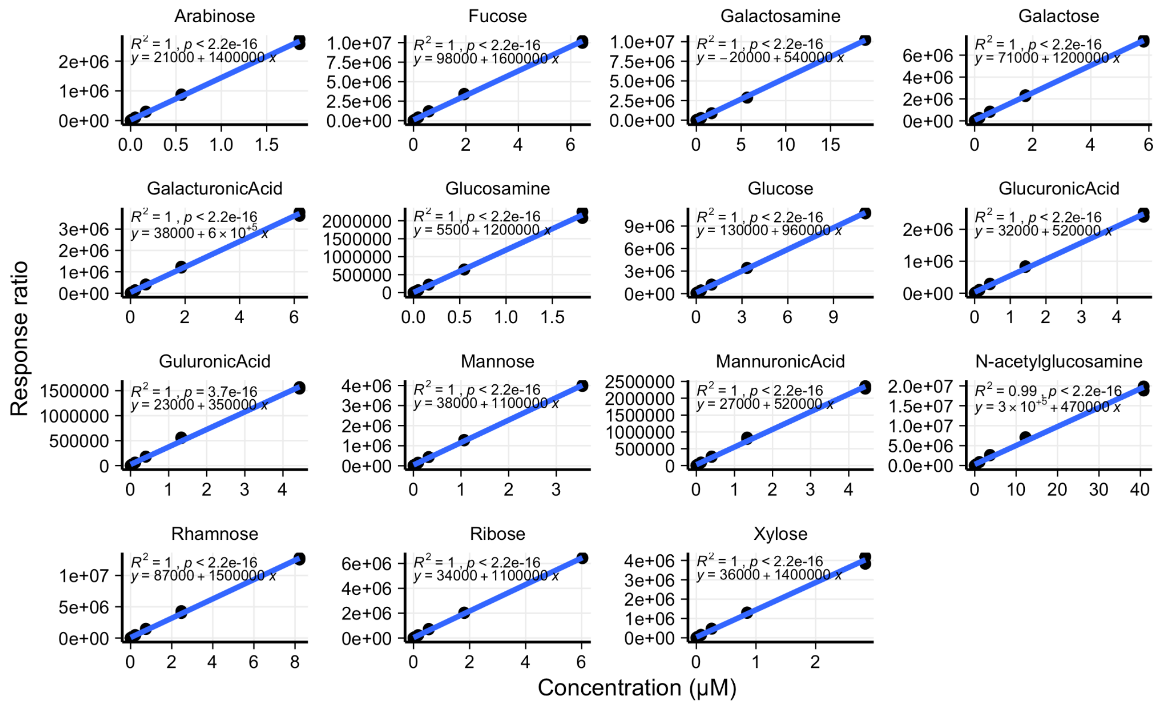


**Figure S6 | Standard curves of monosaccharides.** PMP-derivatives of monosaccharides were quantified by LC-MS/MS in multiple reaction monitoring mode. The blue line represents the linear regression for each standard. Each standard was injected twice, once at the start of the run and once at the end. Response ratios were calculated by normalizing against internal standards.

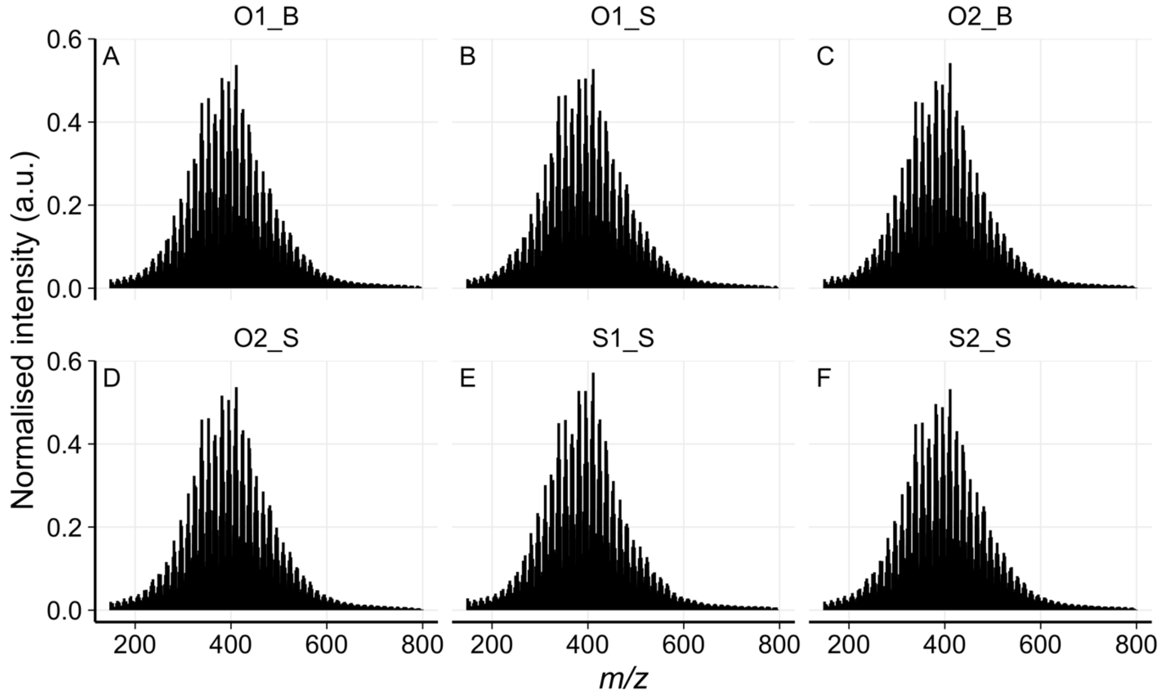


**Figure S7 | FT-ICR-MS analysis of PPL-SPE extracts from MSM107 surface and deep seawater show similar profiles.** Averaged mass spectra of SPE-DOOM seawater extracts show similar Gaussian distributions around 400 *m/z*. Molecular formulae were assigned to 5,263 peaks, with 3,289 shared among all samples that represented 98.12% ± 0.48% (mean ± SD) of the total ion intensity of the samples.

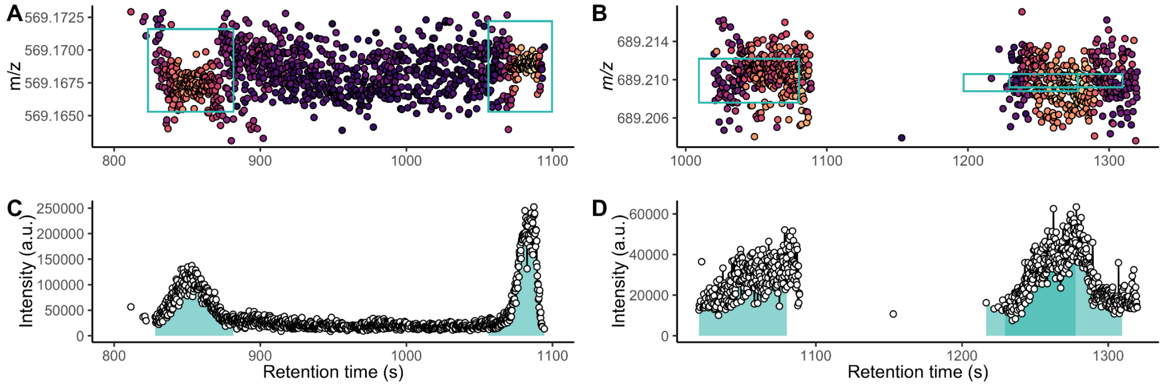


**Figure S8 | Unresolved oligosaccharide isomers hidden within detected features.** Examples of resolved (A, C) and unresolved (B, D) oligosaccharide isomers. A and B show the mass-to-charge chromatograms. Each point represents a scan event and is colored by signal intensity. Feature definitions are depicted by the blue rectangles. C and D show the extracted ion chromatograms. Each point represents a scan event. The blue shading under the curve represents the integrated peak area for the respective features in this sample (EN683-DCM3). A and C represent putative tetrapentose $[M+Na]^+$ or acetylated trihexose $[M+Na]^+$ ions (theoretical *m/z* is 569.1688 for both). B and D represent putative tetrahexose $[M+Na]^+$ ions (theoretical *m/z* is 689.2111).

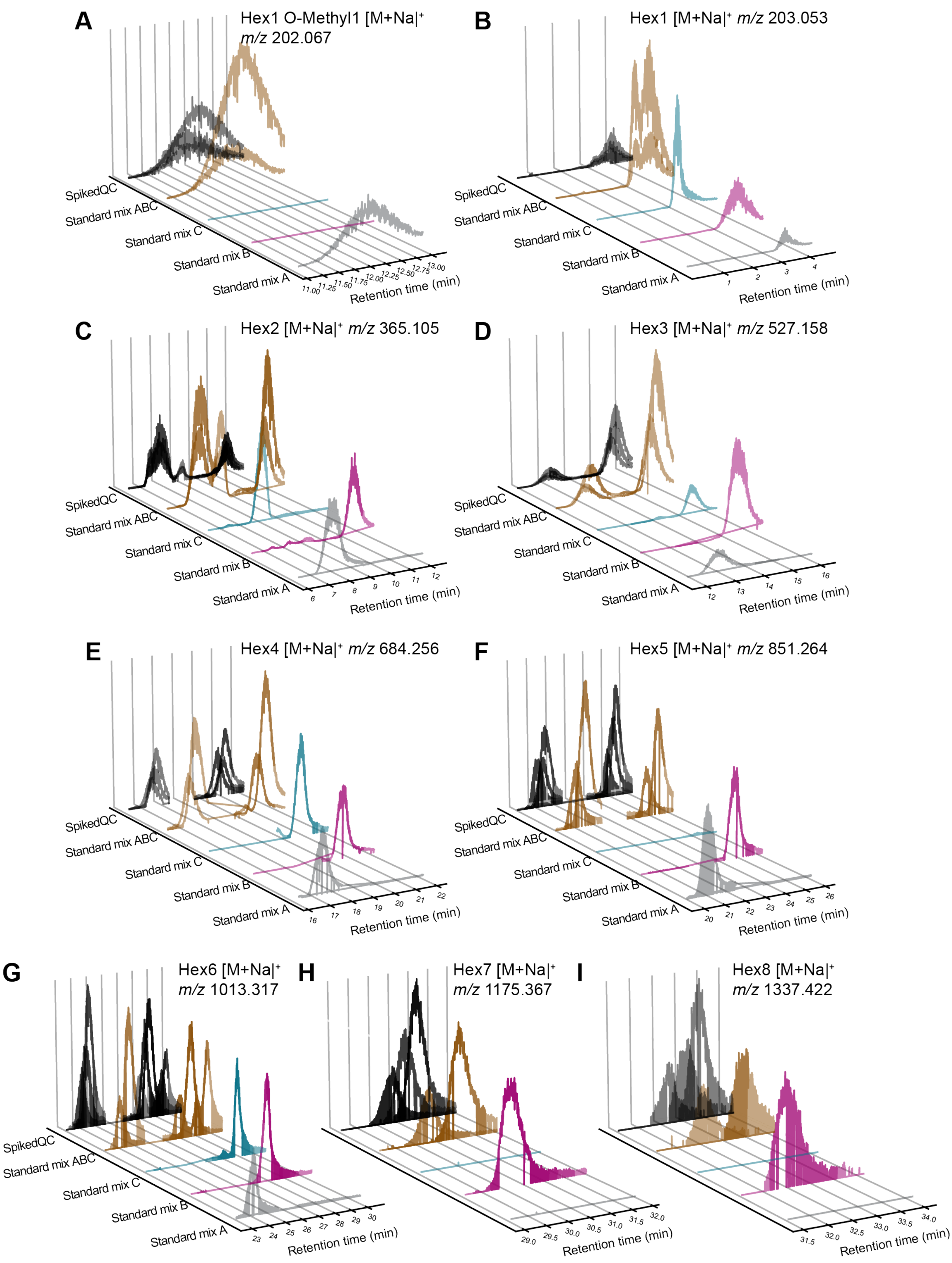


**Figure S9 | Extracted ion chromatograms for standards in separated and combined mixtures and spiked in quality control (QC) samples.** Standard mixtures A (gray line), B (pink line), and C (blue line) are summarized in Table S3. Standard mixture ABC (brown line) is a mixture of all three. Standard mixture ABC was added to the pooled quality controls for the spiked QC samples (black line). Each panel shows the extracted ion chromatograms for a different standard. The minimum and maximum *m/z* and retention time values used are the feature definitions from the pre-processing steps. The annotations from GlycoAnnotateR and the median *m/z* value of the feature are shown in the top right of each panel.

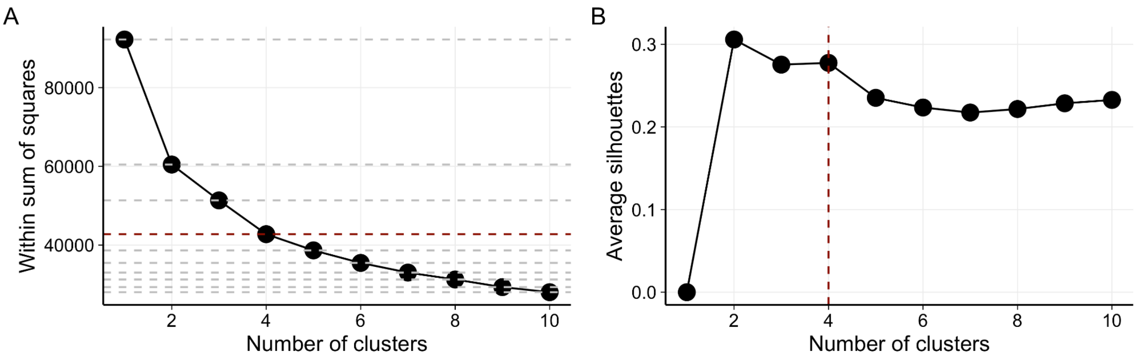


**Figure S10 | Features were grouped into four clusters by *k*-means clustering.** Based on the additional variation (within sum of squares) explained (A) and change in average silhouette score (B) with each additional cluster, *k*=4 was used for the analysis.

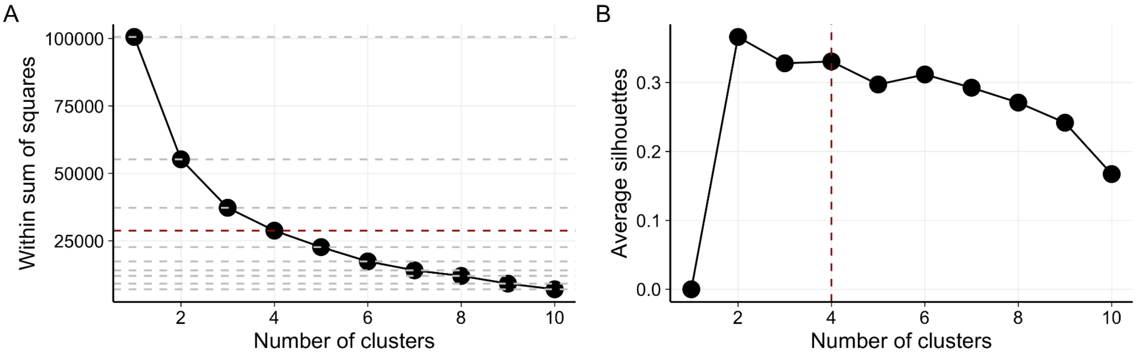


**Figure S11 | Samples were grouped into four clusters by *k*-means clustering**. Based on the additional variation (within sum of squares) explained (A) and change in average silhouette score (B) with each additional cluster, *k*=4 was used for the analysis.

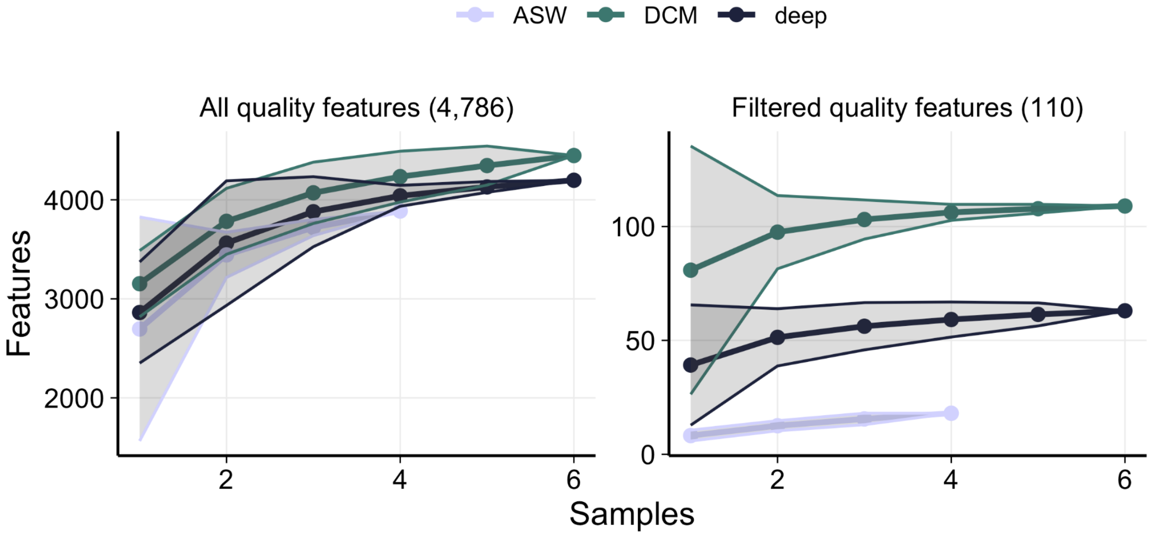


**Figure S12 | Saturation curves suggest that more oligosaccharides are detectable in surface than deep ocean.** Feature accumulation curves generated permutations, with samples added in random order. Features were considered present if integrated signal intensity was >1% of the maximum intensity in that sample. Ranges of permutations are shaded. ASW = artificial seawater; DCM = deep chlorophyll maximum.

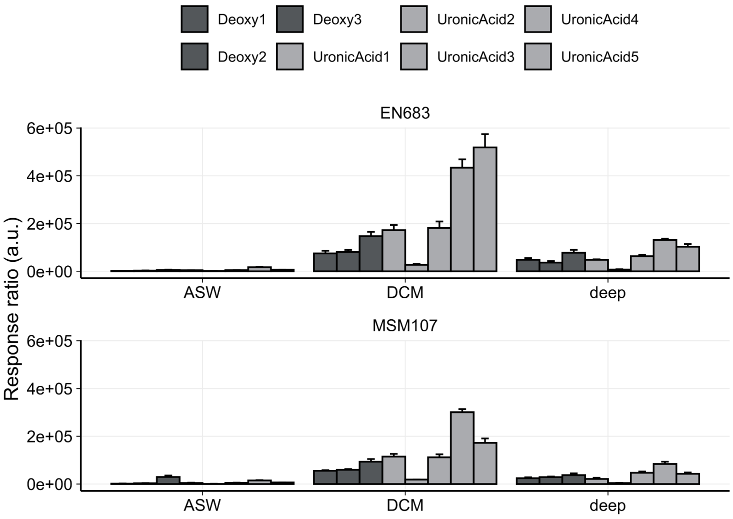


**Figure S13 | Unidentified deoxyhexoses and uronic acids were detected in surface and deep seawater samples.** Three deoxyhexoses and five uronic acids were detected with retention times distinct from standards and therefore could not be identified or quantified. Response ratios shown here correspond to peak areas normalized with respect to isotopically labeled internal standards. Bars shown means and error bars the standard errors of the means.

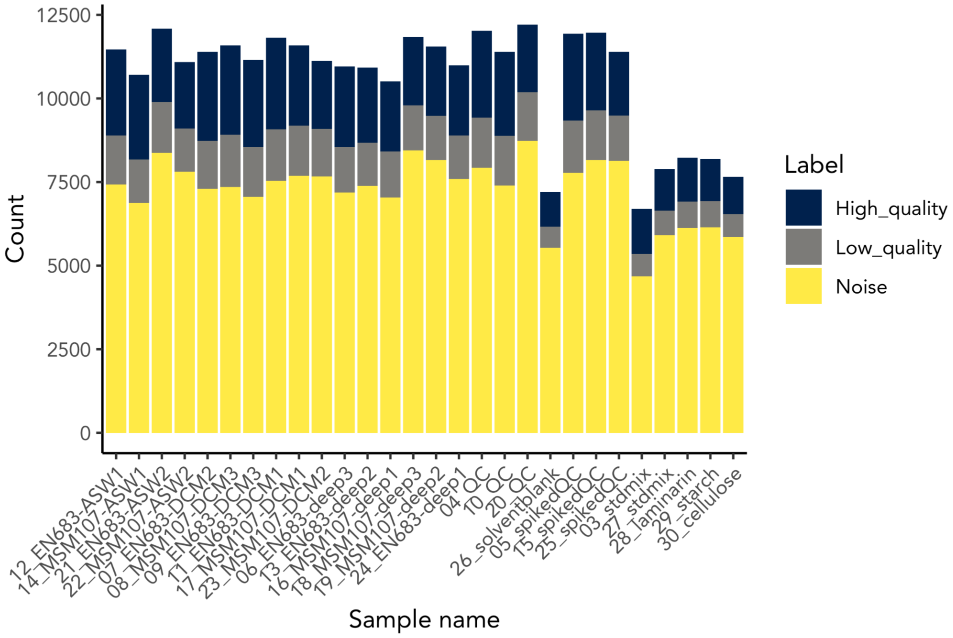


**Figure S14 | The majority of peaks were classified as noise by NeatMS**. Counts of the number of peaks classified as high quality, low quality or noise by NeatMS with our trained model. Features that contained only peaks assigned a noise label were discarded from further analyses.

### Supplementary tables

**Table S1 | Techniques to prepare marine organics for molecular analysis.**

| Technique | Size range targeted | Polarity targeted | Concentration factor | References |
|---|---|---|---|---|
| styrene divinylbenzene solid phase extraction (PPL-SPE) | Indiscriminate, acid induces depolymerization | non-polar | up to x1000 | (2–4) |
| Tangential flow filtration/ ultrafiltration | >1000 Da | Indiscriminate | x100 | (5, 6) |
| Reverse osmosis-electrodialysis | indiscriminate | neutral | x20 | (7–9) |
| Boronate affinity chromatography | Indiscriminate? | cis-diol containing molecules | x20 | (10–12) Unpublished results |
| SeaMet derivatization | >400 Da | indiscriminate | No concentration | (4, 13, 14) |
| Benzoate derivatization followed by PPL-SPE | >550 Da | Extended from non-polar | x50 | (15, 16) |

**Table S2 | Locations and depths sampled for oligosaccharides and dissolved organic matter during two research cruises in the North Atlantic in late May and early June 2022.** Deep chlorophyll maximum (DCM; called surface in this study) and deep ocean seawater were collected in two casts on EN683 (R/V Endeavor), and one cast on MSM107 (R/V Maria S. Merian) for oligosaccharide extraction. SPE-DOM was extracted from seawater collected at four stations on MSM107.

| Location | Coordinates | Expedition and station | Date | Local time | Surface depth (m) | Deep depth (m) | Sample(s) collected |
|---|---|---|---|---|---|---|---|
| Newfoundland | 42.86173°, -53.86997 | EN683, station 19 | 31.05.22 | 06:45 | - | 4,092 | Oligosaccharides |
| Newfoundland | 43.03747°, -54.87668 | EN683, station 19 | 03.06.22 | 11:36 | 125 | - | Oligosaccharides |
| Ireland | 55.106817°, -9.6942° | MSM107, station O2 | 27.05.22 | 16:08 | 35 | 2,300 | Oligosaccharides and SPE-DOM |
| Ireland | 55.2986°, -10.63915° | MSM107, station O1 | 24.05.22 | 06:22 | 30 | 2,500 | SPE-DOM |
| Ireland | 54.946383°, -9.829767° | MSM107, station S1 | 21.05.22 | 21:34 | 30 | - | SPE-DOM |
| Ireland | 55.10695°, -9.69465° | MSM107, station S2 | 29.05.22 | 15:49 | 20 | - | SPE-DOM |

**Table S3 | Sample injection order during LC-HRMS analysis.** Standard mixtures A, B and C are described in Table S4. Quality controls (QC) are a mixture of all samples (i.e., pooled QCs). Spiked QCs are QCs spiked with standard mixtures A, B and C.

| Injection | Name | Sample type | Cruise | Extract type |
|---|---|---|---|---|
| 1 | 03_stdmixABC | standard | | |
| 2 | 04_QC | QC | | |
| 3 | 05_spikedQC | spiked QC | | |
| 4 | 06_EN683-deep3 | deep | EN683 | deep |
| 5 | 07_EN683-DCM2 | DCM | EN683 | DCM |
| 6 | 08_MSM107-DCM3 | DCM | MSM107 | DCM |
| 7 | 09_EN683-DCM3 | DCM | EN683 | DCM |
| 8 | 10_QC | QC | | |
| 9 | 11_EN683-DCM1 | DCM | EN683 | DCM |
| 10 | 12_EN683-ASW1 | ASW | EN683 | ASW |
| 11 | 13_EN683-deep2 | deep | EN683 | deep |
| 12 | 14_MSM107-ASW1 | ASW | MSM107 | ASW |

| | | | | |
|---|---|---|---|---|
| 13 | 15_spikedQC | spiked QC | | |
| 14 | 16_MSM107-deep1 | deep | MSM107 | deep |
| 15 | 17_MSM107-DCM1 | DCM | MSM107 | DCM |
| 16 | 18_MSM107-deep3 | deep | MSM107 | deep |
| 17 | 19_MSM107-deep2 | deep | MSM107 | deep |
| 18 | 20_QC | QC | | |
| 19 | 21_EN683-ASW2 | ASW | EN683 | ASW |
| 20 | 22_MSM107-ASW2 | ASW | MSM107 | ASW |
| 21 | 23_MSM107-DCM2 | DCM | MSM107 | DCM |
| 22 | 24_EN683-deep1 | deep | EN683 | deep |
| 23 | 25_spikedQC | spiked QC | | |
| 24 | 26_solventblank | solvent blank | | |
| 25 | 27_stdmixABC | standard | | |
| 26 | 28_stdmixA | standard | | |
| 27 | 29_stdmixB | standard | | |
| 28 | 30_stdmixC | standard | | |

**Table S4 | Oligosaccharide standard mixtures.**

| Standard mix | Compound | Supplier | Concentration (ng/mL) | Monoisotopic mass (Da) | $[M+Na]^+$ *m/z* |
|---|---|---|---|---|---|
| A | glucose | Sigma Aldrich | 180.06339 | 180.063 | 203.053 |
| A | glucosamine | Sigma Aldrich | 179.079374 | 179.079 | 202.069 |
| A | laminaribiose | Megazyme | 342.116215 | 342.116 | 365.105 |
| A | laminaritriose | Megazyme | 504.16904 | 504.169 | 527.158 |
| A | laminaritetraose | Megazyme | 666.221865 | 666.222 | 689.211 |
| A | laminaripentaose | Megazyme | 828.27469 | 828.275 | 851.264 |
| A | laminarihexaose | Megazyme | 990.327515 | 990.328 | 1013.317 |
| B | galacturonic acid | Sigma Aldrich | 194.042655 | 194.043 | 217.032 |
| B | galactose | Sigma Aldrich | 180.06339 | 180.063 | 203.053 |
| B | 6-O-methyl-galactose | Sigma Aldrich | 194.07904 | 194.079 | 217.068 |
| B | isomaltose | Megazyme | 342.116215 | 342.116 | 365.105 |
| B | maltotriose | Megazyme | 504.16904 | 504.169 | 527.158 |
| B | maltotetraose | Megazyme | 666.221865 | 666.222 | 689.211 |
| B | maltopentaose | Megazyme | 828.27469 | 828.275 | 851.264 |
| B | maltohexaose | Megazyme | 990.327515 | 990.328 | 1013.32 |
| B | maltoheptaose | Megazyme | 1152.38034 | 1152.38 | 1175.37 |
| B | maltooctaose | Megazyme | 1314.43317 | 1314.43 | 1337.423 |
| C | mannose | Sigma Aldrich | 180.06339 | 180.063 | 203.053 |

| | | | | | |
|---|---|---|---|---|---|
| C | mannitol | Sigma Aldrich | 182.07904 | 182.079 | 205.068 |
| C | O-methyl-mannose | Sigma Aldrich | 194.07904 | 194.079 | 217.068 |
| C | fucose | Carbosynth | 164.068475 | 164.068 | 187.058 |
| C | xylose | Sigma Aldrich | 150.052824 | 150.053 | 173.043 |
| C | glucuronic acid | Sigma Aldrich | 194.042655 | 194.043 | 217.032 |
| C | cellobiose | Megazyme | 342.116215 | 342.116 | 365.105 |
| C | cellotriose | Megazyme | 504.16904 | 504.169 | 527.158 |
| C | cellotetraose | Megazyme | 666.221865 | 666.222 | 689.211 |
| C | cellohexaose | Megazyme | 990.327515 | 990.328 | 1013.317 |

**Table S5 | XCMS parameters** (17) **used for pre-processing of LC-HRMS data.** Where possible, parameters were optimized with IPO (18). Default values were used for parameters that are not specified. *Value after retention time correction

| Step and function | Parameter | Value |
|---|---|---|
| Peak picking (findChromPeaks) | min_peakwidth | 3 |
| | max_peakwidth | 130 |
| | ppm | 15.4 |
| | mzdiff | -0.02465 |
| Peak merging (refineChromPeaks) | expandRt | 5 |
| | minProp | 0.5 |
| Peak grouping (groupChromPeaks) and alignment (adjustRtime) | bw | 6 (5*) |
| | minFraction | 2/3 (1/6*) |
| | sampleGroups | sample_type |
| | subset | sample_type == “QC” |
| | span | 0.4 |
| | subsetAdjust | average |

**Table S6 | Metrics of NeatMS model after transfer learning of default model classifier over 10,000 epochs.**

| Metric | Value |
|---|---|
| Threshold | 0.16 |
| True positive rate | 1.00 |
| False positive rate | 0.09 |
| Area under ROC | 0.98 |

**Table S7 | Quantification parameters for monosaccharides.** Retention times for each of the PMP-derivatized monosaccharides that were quantified with the specified multiple reaction monitoring (MRM) transitions. Calibration curves were calculated after normalization with respect to internal standards.

| Compound | bis-PMP mass (Da) | Precursor ion (m/z) | Product ion (m/z) | Collision energy (eV) | Retention time (min) | Calibration curve | $R^2$ |
|---|---|---|---|---|---|---|---|
| Ribose | 480.2 | 481.2 | 175.1 | 35 | 3.65 | y = 1067336x + 33847 | 0.999 |
| Arabinose | 480.2 | 481.2 | 175.1 | 35 | 6.1 | y = 1424178x + 21163 | 0.998 |
| Xylose | 480.2 | 481.2 | 175.1 | 35 | 6.23 | y = 1409923x + 36232 | 0.997 |
| Rhamnose | 494.2 | 495.2 | 175.1 | 35 | 3.86 | y = 1543114x + 86860 | 0.999 |
| Fucose | 494.2 | 495.2 | 175.1 | 35 | 6.83 | y = 1567434x + 98197 | 0.998 |
| Galactosamine | 509.2 | 510.2 | 175.1 | 35 | 3.25 | y = 540622x + -19595 | 1.000 |
| Glucosamine | 509.2 | 510.2 | 175.1 | 35 | 2.04 | y = 1182281x + 5536 | 0.998 |
| Mannose | 510.2 | 511.2 | 175.1 | 35 | 2.72 | y = 1119725x + 38410 | 1.000 |
| Glucose | 510.2 | 511.2 | 175.1 | 35 | 5.52 | y = 960866x + 125183 | 1.000 |
| Galactose | 510.2 | 511.2 | 175.1 | 35 | 5.66 | y = 1248510x + 70999 | 0.999 |
| MannuronicAcid | 524.2 | 525.2 | 175.1 | 35 | 2.85 | y = 523855x + 27175 | 0.997 |
| GuluronicAcid | 524.2 | 525.2 | 175.1 | 35 | 3.45 | y = 350628x + 23212 | 0.997 |
| GlucuronicAcid | 524.2 | 525.2 | 175.1 | 35 | 5.38 | y = 515783x + 32220 | 0.997 |
| GalacturonicAcid | 524.2 | 525.2 | 175.1 | 35 | 5.57 | y = 595179x + 37912 | 0.998 |
| N-acetylglucosamine | 551.2 | 552.2 | 175.1 | 35 | 5.1 | y = 474540x + 296193 | 0.994 |

**Table S8 | Dissolved organic carbon (DOC) concentrations in PPL-SPE extracts.** Values represent DOC concentration of SPE-DOM in seawater.

| Station | Depth | Replicate | [DOC] (µM) |
|---|---|---|---|
| O1 | Bottom | 1 | 39.4 |
| | | 2 | 40.8 |
| | | 3 | 13.8 |
| | Surface | 1 | 46.7 |
| | | 2 | 48.2 |

| | | | |
|---|---|---|---|
| | | 3 | 46.7 |
| O2 | Bottom | 1 | 40.4 |
| | | 2 | 13.8 |
| | | 3 | 38.5 |
| | Surface | 1 | 17.9 |
| | | 2 | 46.9 |
| | | 3 | 45.5 |
| S1 | Surface | 1 | 49.2 |
| | | 2 | 30.5 |
| | | 3 | 47.4 |
| S2 | Surface | 1 | 49.0 |
| | | 2 | 48.9 |
| | | 3 | 20.8 |

**Table S9 | Molecular indices calculated from FT-ICR mass spectrometer analysis of SPE-DOM composition.** IDEG is a degradation index that reflects how refractory the DOM is in a sample (19, 20). IBIO indicates the extent of recent biological production (21). MLB is a measure of the saturation of a sample (22). CRAM is a measure of refractory DOM components (5).

| Station | Water layer | $I_{DEG}$ | $I_{BIO}$ | MLB | CRAM (%) |
|---|---|---|---|---|---|
| S1 | Surface | 0.7042 | 0.3439 | 5.647 | 69.09 |
| S2 | Surface | 0.7053 | 0.3393 | 5.73 | 68.75 |
| O1 | Surface | 0.6914 | 0.3462 | 6.037 | 69.27 |
| O2 | Surface | 0.7171 | 0.3287 | 5.398 | 69.58 |
| O1 | Bottom | 0.7086 | 0.3395 | 5.664 | 69.06 |
| O2 | Bottom | 0.6933 | 0.351 | 5.938 | 68.09 |

**Table S10 | Literature values for fucose enrichment at depth. HMW = high molecular weight.**

| Reference | Fraction | Location | Fucose at surface (%) | Fucose at depth (%) | Fold enrichment | Monomers quantified | Quantification method |
|---|---|---|---|---|---|---|---|
| (23) | Combined | Equatorial Pacific | 18 | 26 | 1.44 | Neutral | HPLC-PAD |
| (24) | HMW | North Pacific Subtropical Gyre | 17 | 25 | 1.47 | Neutral | GC-MS of alditol acetates |
| (25) | HMW | North Pacific | 16.4 | 15.6 | 0.95 | Neutral | GC-MS of alditol acetates |
| (25) | HMW | Gulf of Mexico | 15.1 | 18.8 | 1.24 | Neutral | GC-MS of alditol acetates |
| (25) | HMW | Sargasso Sea | 17.3 | 18.5 | 1.07 | Neutral | GC-MS of alditol acetates |
| (26) | HMW | Pacific (2°S, 140°W) | 14 | 18 | 1.29 | Neutral | HPLC-PAD |
| (26) | HMW | Pacific (12°S, 135°W) | 14 | 13 | 0.93 | Neutral | HPLC-PAD |
| (27) | HMW | Mid-Atlantic Bight | 17 | 20 | 1.18 | Neutral | GC-MS of alditol acetates |

### Extended methods

To understand carbohydrate cycling and longevity in marine environments, this study focused on obtaining molecular-level insights into oligosaccharides dissolved in surface and deep seawater. Environmental samples were collected during two research cruises from productive surface ocean and deep ocean water. Intact oligosaccharides were extracted from seawater and concentrated. Quantification of monosaccharide constituents post-hydrolysis validated the extraction process and enabled compositional and quantitative analysis. Analysis by liquid chromatography high resolution mass spectrometry complemented by a deep learning-based denoising approach and screening for carbohydrates with de novo annotation allowed the detection and differentiation of intact oligosaccharides. This approach aimed at to provide the resolution to uncover the fundamental principles that govern marine carbohydrate turnover and resilience against microbial degradation.

This extended materials and methods section contains detailed descriptions of:

- Physical parameters of sampled water masses
- GGC-SPE of oligosaccharides
- LC-HRMS procedure
- LC-HRMS data analysis, including NeatMS training
- Monosaccharide quantification
- SPE-DOM analysis

**Physical parameters of water masses**

Surface seawater near Newfoundland had a neutral density of 26.604 kg $m^{-1}$ indicative of mixing with Western North Atlantic Central Water (**Fig. 1C**) (28). A neutral density of 26.975 kg $m^{-1}$ of surface seawater sampled near Ireland suggests mixing with Eastern North Atlantic Central Water. Physical parameters of deep ocean seawater sampled from a depth of approximately 4,100 m off Newfoundland with a potential temperature of 1.96°C and salinity of 34.89 aligned with North Atlantic Deep Water influenced by Danish Straits Overflow Water (29) (see (30, 31) for comparison). The deep ocean water mass sampled at 2,300 m near Ireland had a higher temperature (2.98°C) and density (35.107 g $kg^{-1}$) suggesting that these samples comprised North Atlantic Deep Water influenced by Iceland-Scotland Overflow water (28). These physical properties indicate that deep ocean water masses sampled near Newfoundland and Ireland represented North Atlantic Deep Water formed in two distinct polar regions.

**Solid-phase extraction of oligosaccharides in detail**

Oligosaccharides were extracted from 600 mL of 0.22 µm filtered seawater in triplicate immediately after sampling and from 600 mL 0.22 µm filtered artificial seawater prepared using 35 g $L^{-1}$ sea salts (Sigma) in duplicate. Oligosaccharides were purified by adapting a non-porous graphitized carbon cartridge solid-phase extraction (GCC-SPE) protocol from (1) using only gravity flow. Cartridges (Supelclean™ ENVI-Carb™ SPE Tube, 60 mL, 10 g) were pre-conditioned with 60 mL acetonitrile followed by 60 mL MilliQ-water before application of a sample. To accommodate the larger sample volumes required by the environmental context, cartridge volumes were extended to 300 mL with 3D-printed containers (**Fig. S1**). After sample application and four 30 mL washes with MilliQ-water, oligosaccharides were eluted three times with 15 mL elution buffer (49.95: 49.95: 0.1 (v/v) MilliQ-water: acetonitrile: formic acid). Elution fractions were collected together in 50 mL tubes and stored at -20°C until further processing on land. 10 mL of eluted oligosaccharides were partially dried by vacuum centrifugation for 4 hours at room temperature in an Eppendorf ® Concentrator Plus (V-HV) to evaporate volatile organics. The remaining volume was freeze-dried (Labogene, ScanVac CoolSafe). After reconstituting the dried samples in 2 mL MilliQ-water, 500 µL were taken for acid hydrolysis and monosaccharide analysis, and the remainder dialyzed to remove any residual salts by using a 100-500 Da membrane (SpectraPor® Biotech) against 15 L of MilliQ-water for 4 weeks, with water changes every 3-5 days. Dialyzed samples were freeze-dried and resuspended in 1 mL MilliQ-water. Half of the volume was dried again by vacuum centrifugation and resuspended in 30 µL of MilliQ-water for analysis with liquid chromatography high resolution mass spectrometry (LC-HRMS).

**Liquid chromatography high resolution mass spectrometry (LC-HRMS)**

Of each sample, 5 µL were taken and combined to generate a quality control standard (QC). The QC was split in half (40 µL each), and one half was spiked with mono- and oligosaccharide standards (mixes A, B and C, see **Table S4**). Acetonitrile was added to all QC and seawater samples for a final concentration (v/v) of 70%. The approximate final concentration factor of oligosaccharides relative to seawater was 400x. Samples were analyzed by LC-HRMS in random order with repeated runs of QCs every 5 injections (**Table S3**).

LC-HRMS measurements were performed on a Thermo Fisher Vanquish Horizon UHPLC coupled to a Thermo Fisher Q-Exactive Plus MS. The system was controlled using Chromeleon software (version 7.2.10; Thermo Fisher Scientific, Waltham, USA). 10 µL of samples were injected for separation on a Accucore-150-amide HILIC column with the dimensions 100 mm long, 2.1 mm diameter and a particle size of 2.6 µm (Thermo Scientific, USA). For injection order see **Table S3**. Mobile phase solvents consisted of 10 mM ammonium formate at pH 5 for solvent A and UHPLC-grade acetonitrile for solvent B. During a run at a flow rate of 0.4 mL $min^{-1}$, solvent B was held at 90% for 1 min after injection and then gradually decreased from 90% to 60% B over 40 min. After the gradient, B was further reduced to 40% over 4 min, increased back to 90% over a 3 min period and kept at 90% B for 16 min to recondition the column. Oligosaccharides in samples were detected with the following MS conditions: positive ionization mode with a spray voltage of 3.9 kV; sheath, auxiliary and sweep gas flow rates, 50, 13 and 3 respectively; capillary temperature, 300°C; S-lens RF level, 60; auxiliary gas heater temperature, 320°C; mass resolution, 70,000 at *m/z* 200; scan range, *m/z* 175 to 1,400.

**LC-HRMS data analysis in detail**

LC-HRMS data were pre-processed by picking and refining peaks, aligning retention times and grouping peaks across samples to obtain features defined in *m/z* and retention time dimensions. Raw files from Thermo Fisher were first converted to mzML format using the ProteoWizard command line tool 'msconvert' version 3.0.20239 with a 'peakPicking' filter. Pre-processing (peak picking, peak merging, retention time alignment, peak grouping and peak filling) steps were performed using the ‘XCMS’ package (v4.0.2) in R (17). Peak picking and retention time alignment parameters were optimized using IPO (v1.20.0) (18). Non-default parameters for each pre-processing step are provided in **Table S5**. CAMERA (v1.56.0) was used to group peaks again by retention time using peak full-width half-maximum values for group borders (perfFWHM = 0.6), which enabled detection of isotopes (ppm = 3) (32). Features annotated as $^{13}C$-isotopes were removed.

We trained and applied the deep learning-based peak filter tool NeatMS (33) to classify peaks within the features into high and low quality and noise. 2000 peaks from 9 samples (three spiked QCs, three QCs, one deep ocean, one surface and one processing blank) were manually classified and reviewed (**Fig S3**). A neural network handler was created (matrice size = 120, margin = 1, min_scan_num = 5) and used to split the classified peaks into batches (80% training, 10% test, 10% validation) with no normalization of classes. Transfer learning trained the classifier of the default NeatMS model for 10,000 epochs at a learning rate set to 0.000001 with "Adam" as optimizer using the 'get_threshold' method to compute the optimal threshold (github.com/bihealth/NeatMS/blob/master/data/model/neatms_default_model.h5). The test data set had an accuracy of 81.71% and loss of 0.4448 (**Fig. S4**). Model metrics are given in **Table S6**.

Applying the trained model to the dataset of LC-MS features classified 70,270 of 270,061 peaks as high or low quality for further assessment (**Fig S14**). The high proportion of peaks assigned as noise (~75%) was expected, as our analysis strategy was to start very broad and successively narrow down the signals retained. In this way we could be more confident that no true signals were lost, especially during peak picking, but in this trade-off, we also have more noise in our data at the start of the pipeline. Features with retention times before 1 min or after 35 min were also discarded, leaving 33,433 peaks in 8,408 features across all samples. Seawater extract samples comprised 21,830 peaks in 4,786 features.

We searched the quality features for putative oligosaccharides by comparing *m/z* values to theoretical oligosaccharides. Theoretical oligosaccharide formulae, masses and *m/z* values (H, Na, K and NH4 adducts, all singly charged) were predicted with the R package 'GlycoAnnotateR' (https://github.com/margotbligh/GlycoAnnotateR). Ionization type was set to 'ESI' and ionization mode to positive. Allowed modifications were carboxylic acid, deoxy, N-acetyl, O-acetyl, O-methyl, amino, anhydro-bridge, alditol and sulfate, for pentoses and hexoses with degrees of polymerisation (DP) 2-6. For each theoretical oligosaccharide ion, an *m/z* interval was created as *m/z* ± 3 ppm. Annotations were assigned if the median *m/z* value of a feature was within the interval. Further filtering was applied to remove features with >60% of the maximum integrated intensity for that feature in any of the processing blanks (34), and if the integrated intensity for any peak in a feature was greater than 1E8 (**Fig. S5**). The 555 peaks of 189 features that matched theoretical oligosaccharides in *m/z* and were not abundant in processing blanks were manually re-classified as quality and noise. This step was necessary due to the relatively high false positive rate (0.09) at the model threshold. Again, we purposefully chose a model threshold with a high true positive rate (1.0) to ensure no true signals were lost, but the consequence of this was a high false positive rate. Manual classification returned 110 quality features.

**Monosaccharide analysis in detail**

For acid hydrolysis prior to monosaccharide quantification, aliquots of 500 µL were transferred to pre-combusted (450°C, 4.5h) glass ampoules and combined with 500 µL of 2 M HCl (24 h, 100°C). After hydrolysis, acid and water were evaporated in an acid-proof vacuum concentrator (Martin Christ Gefriertrocknungsanlagen GmbH, Germany) and dried samples reconstituted in 500 µL MilliQ-water.

Following a previously published protocol, acid hydrolyzed samples containing monosaccharides (25 µL) were derivatized with 75 µL of 0.1M 1-phenyl-3-methyl-5-pyrazolone (PMP) in 2:1 methanol:ddH2O with 0.4 % ammonium hydroxide for 100 minutes at 70°C (35). Additionally, each sample was spiked with an internal standard of 10 µM $^{13}C6$-Glucose, $^{13}C6$-Galactose and $^{13}C6$-Mannose (mass 186 Da). For quantification, we derivatized a serial dilution of a standard mix containing Galacturonic acid, D-Glucuronic acid, Mannuronic Acid, Guluronic Acid, Xylose, Arabinose, D-Glucosamine, Fucose, Glucose, Galactose, Mannose, N-Acetyl-D-glucosamine, N-Acetyl-D-galactosamine, N-Acetyl-D-mannosamine, Ribose, Rhamnose and D-galactosamine. Samples and standards were derivatized by incubation at 70°C for 100 minutes. After derivatization, samples were diluted 1:100 in 0.1% formic acid.

Following (36), PMP-derivatives were measured on a SCIEX qTRAP5500 and an Agilent 1290 Infinity II LC system equipped with a Waters CORTECS UPLC C18 Column, 90Å, 1.6 µm, 2.1 mm X 50 mm reversed phase column with guard column. The mobile phase consisted of buffer A (10 mM NH4Formate in ddH2O, 0.1% formic acid) and buffer B (100% acetonitrile, 0.1% formic acid). PMP-derivatives were separated with an initial isocratic flow of 15% Buffer B for 2 minutes, followed by a gradient from 15 % to 20% Buffer B over 5 minutes at a constant flow rate of 0.5 ml/min and a column temperature of 50°C. The ESI source settings were 625°C, with curtain gas set to 30 (arbitrary units), collision gas to medium, ion spray voltage 5500 (arbitrary units), temperature to 625°C, Ion source Gas 1 &2 to 90 (arbitrary units). PMP-derivatives were measured by multiple reaction monitoring (MRM) in positive mode with previously optimized transitions and collision energies. For example, a glucose derivative has a Q1 mass of 511 and was fragmented with a collision energy of 35V to yield the quantifier ion of 175Da and the diagnostic fragment of 217Da. Different PMP-derivatives were identified by their mass and retention in comparison to known standards. Technical variation in sample processing were normalized by the amount of internal standard in each sample. Peak areas of the 175 Da fragment were used for quantification using an external standard ranging from 100 pM to 10 µM (**Table S7, Fig S6**). Raw files from Sciex (wiff format) were first to mzML format using the ProteoWizard command line tool 'msconvert' version 3.0.20239 with a 'peakPicking' filter for MS levels 1 and 2, and chromatograms were imported to R with the ‘readSRMdata’ from MSnbase (37).

**SPE-DOM and FT-ICR-MS in detail**

Filtered seawater samples (2 L each) collected in triplicate at four stations northwest of Ireland (**Table S2, Fig. 1B**) were acidified to pH 2 and dissolved organic matter was solid phase-extracted using styrene divinylbenzene polymer cartridges (PPL) (2). Analysis of this SPE-DOM consisted of direct injection mass spectrometry with technical duplicates on a Bruker Solarix XR 15 Tesla Fourier transform ion cyclotron resonance mass spectrometer (Bruker Daltonik GmbH, Bremen, Germany), equipped with a 15 Tesla superconducting magnet (Bruker Biospin, Wissembourg, France) and an electrospray ionization source (Apollo II ion source, Bruker Daltonik GmbH) in negative ionization mode at 4.5 kV (38). Samples were introduced into the microelectrospray source (Agilent, Waldbronn, Germany) at a flow rate of 120 µl $h^{-1}$ with a nebulizer gas pressure of 20 psi and a drying gas pressure of 15 psi (250°C). 2-3 technical replicates were injected for each sample. For analysis validation, an in-house reference sample (North Equatorial Pacific Intermediate Water (NEqPIW); www.icbm.de\en\ds-dom), collected at the Natural Energy Laboratory of Hawaii Authority in 2009, was measured twice per day (39). FT-ICR mass spectra with *m/z* from 150 to 2000 were exported to peak lists after internal calibration in Bruker Daltonik Data Analysis software (calibrated to errors <0.02 ppm). Calibrated mass spectra were further processed, and molecular formulae assigned to peaks using the ICBM-OCEAN processing pipeline (40). The data include the likeliest chemical formula for a given mass chosen based on the most isotope ratio verifications, the greatest homologous series network, and the smallest difference to the reference mass.

The output of ICBM-OCEAN was then further processed in R. Spectra were filtered to only include peaks with *m/z* values between 150 and 800. Peaks assigned molecular formulae with isotopologues or that contained P were also removed. Only peaks with molecular formulae with O/C ratios between 0.05 and 1, and H/C ratios below 1.8 were retained in a further filtering step. Any peaks present in the blanks were additionally removed from the data. Intensity values were then averaged across technical (injection) replicates. Peaks with values below the method detection limit were then set to have zero intensity values. Intensity values were then averaged across extraction triplicates. Finally, intensity values were normalized to the total ion intensity of each sample.

Molecular indices were then calculated based on the processed, averaged and normalized spectra. The results are summarized in **Table S8**. $I_{DEG}$ was calculated as previously described (20). Briefly, the sum of the intensities of negatively correlating compounds ($C_{21}H_{26}O_{11}$, $C_{17}H_{20}O_9$, $C_{19}H_{22}O_{10}$, $C_{20}H_{22}O_{10}$, $C_{20}H_{24}O_{11}$) was divided by the sum of negatively correlating and positively correlating compounds ($C_{13}H_{18}O_7$, $C_{14}H_{20}O_7$, $C_{15}H_{22}O_7$, $C_{15}H_{22}O_8$, $C_{16}H_{24}O_8$). $I_{BIO}$ was calculated as previously described (21) as the ratio between the sum of the intensities of $C_{13}H_{18}O_5$, $C_{13}H_{16}O_6$, $C1_3H_{19}NO_6$, $C_{13}H_{17}NO_7$, $C1_8H_{28}O_7$ and the sum of the intensities of $C_{14}H_{16}O_8$, $C_{17}H_{24}O_8$, $C_{16}H_{22}O_9$, $C_{19}H_{28}O_9$, $C_{19}H_{28}O_{10}$. MLB (molecular lability boundary) was calculated as described by (22) as the sum of the intensities of peaks with H/C ratios greater than or equal to 1.5. The percentage of CRAM was calculated as the sum of the intensities of CRAM-like signals as defined by ratios of elements and double bond equivalents (DBE) (5).

The global ocean FT-ICR-MS data (41) were processed in the same way as the samples in this study. Potential temperature ($\theta$) and potential density anomaly ($\sigma_\theta$) were calculated using ‘oce’ (42), and from those values apparent oxygen utilization (AOU) was calculated. The water masses were defined as summarized in the table below, and values were averaged for each water mass:

| Water mass | Ocean | Latitude | Potential temperature | Temperature (°C) | Salinity (PSU) | Depth (m) | AOU |
|---|---|---|---|---|---|---|---|
| SSW | Atlantic | 20-40 | >15 | - | - | <50 | - |
| NADW | Atlantic | >20 | >2 | <4 | >34.8 | >1000 | - |
| AABW | Atlantic | <-50 | <2 | - | >34.6 | >2000 | - |
| EqSW | Atlantic | -20-20 | >15 | - | - | <50 | - |
| AASW | Atlantic | <-50 | <2 | - | - | <50 | - |
| CDW | Pacific | 0-40 | <2 | - | >34.6 | >2000 | - |
| PDW | Pacific | >40 | - | - | - | 500-2000 | <50 |
| AAIW_Atl | Atlantic | <50 | <4 | - | <34.5 | 600-1400 | - |
| AAIW_Pac | Pacific | <50 | <4 | - | <34-5 | 600-1400 | - |
| NPIW | Pacific | >40 | - | - | <34.5 | <600 | <40 |

SSW = Subtropical Surface Water; NADW = North Atlantic Deep Water; AABW = Antarctic Bottom Water; EqSW = Equatorial Surface Water; AASW = Antarctic Surface Water; CDW = Circumpolar Deep Water; PDW = Pacific Deep Water; AAIW_Atl = Antarctic Intermediate Water (Atlantic); AAIW_Pac = Antarctic Intermediate Water (Pacific); NPIW = North Pacific Intermediate Water.
